# Radius-Optimized Efficient Template Matching for Lesion Detection from Brain Images

**DOI:** 10.1101/2020.01.10.893099

**Authors:** Subhranil Koley, Pranab K. Dutta, Iman Aganj

## Abstract

Computer-aided detection of brain lesions from volumetric magnetic resonance imaging (MRI) is in demand for fast and automatic diagnosis of neural diseases. The template-matching technique can provide satisfactory outcome for automatic localization of brain lesions; however, finding the optimal template size that maximizes similarity of the template and the lesion remains challenging. This increases the complexity of the algorithm and the requirement for computational resources, while processing large MRI volumes with three-dimensional (3D) templates. Hence, reducing the computational complexity of template matching is needed. In this paper, we first propose a mathematical framework for computing the normalized cross-correlation coefficient (NCCC) as the similarity measure between the MRI volume and approximated 3D Gaussian template with linear time complexity, 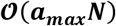, as opposed to the conventional fast Fourier transform (FFT) based approach with the complexity 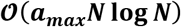, where ***N*** is the number of voxels in the image and ***a**_max_* is the number of tried template radii. We then propose a mathematical formulation to analytically estimate the optimal template radius for each voxel in the image and compute the NCCC with the location-dependent optimal radius, reducing the complexity to 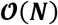. We test our methods on one synthetic and two real multiple-sclerosis databases, and compare their performances in lesion detection with FFT and a state-of-the-art lesion prediction algorithm. We demonstrate through our experiments the efficiency of the proposed methods for brain lesion detection and their comparable performance with existing techniques.

## Introduction

A lesion in the brain represents a diseased area or one with abnormal tissue due to many factors such as infection, traumatic brain injury, tumors, multiple sclerosis (MS), stroke, etc.^1^ The complications resulting from brain lesions are numerous including neurodegeneration, memory or vision loss, and behavioral changes^2^ The presence of a lesion in the brain often demands prompt clinical investigation to initiate the treatment process. Magnetic resonance imaging (MRI) contributes to the diagnosis significantly by visualizing the neuroanatomy with high soft-tissue contrast *in vivo*. Detection of the exact location of lesion is essential for clinical implications and thus assists in diagnosis and treatment planning. Manual assessment (i.e. visual screening) of individual two-dimensional (2D) slices from the whole MRI volume is the conventional practice of identifying the presence of lesions. This is a cumbersome task, especially in the presence of multiple lesions (e.g., MS, metastasis, etc.) or when performing it on a large dataset. Inter- and intra-rater variabilities are other caveats of manual lesion detection. The limitations of manual screening and volumetric analysis by trained experts have led to the development of semi-automatic and automatic segmentation algorithms in the last two decades, which can assist the clinician in identifying the region of interest (ROI) from the magnetic resonance (MR) image. The study conducted by Egger et al.^3^ on fluid attenuation inversion recovery (FLAIR) MRI of MS lesions over 50 cases, reported that the segmentation methods such as lesion growth^4^ and lesion prediction algorithm (LPA)^5^ underperformed for quantifying the number of lesions on the subjects with high lesion load. According to them, the difficulties in the evaluation of the confluent lesions in periventricular region might be the cause of poor performance, which emphasizes the importance of prior detection of lesions in MRI volume, especially when multiple lesions are present in brain.

Machine learning, especially deep learning, has been successfully applied for automatic segmentation of brain lesions.^6^ Deep neural networks are popular because of their high performance over conventional machine-learning techniques. For instance, Lu et al.^7^ adopted transfer learning using AlexNet convolutional architecture to determine the pathological brain slices (images) of various diseases, such as Glioma, MS, Alzheimer’s, Huntington’s, etc., from a T2 MR image database. They employed one layer from each of the fully connected, softmax, and classification layers as the replacement for the last three layers in the original AlexNet, and with this modified network, achieved cent percentage accuracy for detection of the pathological brain from normal slices. In the following year, Lu et al.^8^ proposed a deep neural architecture comprising different steps, such as: batch normalization to improve pre-trained AlexNet, training on normal and abnormal brain MR images, and implementation of extreme learning machine classifier to replace several layers at the end of the network (in which chaotic bat algorithm was used to optimize the number of end layers for replacement). This methodology outperformed different variants of AlexNet with comprehensive accuracy for binary classification of abnormal MR images from healthy samples. The present-day deep learning requires a large amount of training data as well as high computational power. In the context of MRI, different scanners use different acquisition parameters leading to variations in image characteristics. This might create difficulty for supervised approaches, for instance when testing on a previously unseen image that deviates from the training data. The key advantage of using unsupervised approaches is that they do not require any prior knowledge or training data. Cabezas et al.^9^ developed a two-stage unsupervised method consisting of modified expectation-maximization (EM) for initial soft-tissue segmentation from T1-weighted (T1), T2-weighted (T2), and proton-density (PD) images and thresholding of FLAIR images for hyperintense-lesion segmentation. They reported 55% accuracy as the best mean sensitivity obtained in hyperintense lesion detection. Generally, thresholding-based approaches depend on intensity histogram to find the cutoff intensity, based on which the target lesion (hyper- or hypo-intense) could be differentiated. This technique faces difficulties in classifying non-lesion regions having similar intensity profile as lesions, where voxel intensity values are influenced by intensity inhomogeneity or artifacts.^10^ *Template matching*, a fully unsupervised technique, has been employed for automatic lesion detection from medical images because of its satisfactory performance in localization of lesions. A general choice for the template could be a radially-symmetric shape such as a circle in 2D or a sphere in 3D, where the template with varying radii is matched at each image voxel. In other words, the image is locally compared with the template by some similarity measure, and this operation is repeated for a set of radii until the optimal radius is found.^11,12^ Tourassi et al.^13^ developed a template-matching technique with a mutual-information similarity measure to detect mass from digital mammograms. Later on, Lochanambal et al.^14^ defined the template based on the shape and the brightness of target lesion and micro calcifications for lesion detection in mammographic images. Template matching has also been employed for detection of lung nodules; for instance, Farag et al.^15^ proposed deformable 2D and 3D templates for spiral CT scan images, Osman et al.^16^ developed a 3D template to perform convolution with 3D ROI image for serial CT images, and Wang et al.^17^ proposed a 3D spherical template with varying size to detect lesions having a diameter of 4-20 mm for typical CT scan images. Later, Moltz et al.^18^ developed a generalized method based on template matching for detection of various types of lesions, such as lung nodules, liver metastasis, and lymph nodes, from CT images. Next, we discuss the applications of template matching in MRI.

Warfield et al.^19^ developed an “adaptive, template moderated, spatially varying statistical classification” framework for segmentation of brain anatomy of neonates, MS lesions, brain tumors, and damaged knee cartilage from MR images by using an anatomical template to moderate the statistical classification. Ambrosini et al.^10^ proposed a 3D spherical template and used the normalized cross-correlation coefficient (NCCC) as the similarity measure between templates (with varying radii) and the brain volume for automatic detection of metastatic brain tumor from post-contrast T1 MRI (spoiled gradient echo pulse sequence). Later, Farjam et al.^20^ used 3D post-Gd-DTPA T1 volumes to localize hyperintense metastatic lesions with a 3D spherical-shell template. In a different study, Yang et al.^21^ considered contrast-enhanced MRI black-blood pulse sequence for metastatic lesion detection using 3D template. Wang et al.^22^ performed another study with an adaptive template using NCCC as similarity measure for tumor detection using T1 MR images. We found that an exhaustive search was a common choice to find the optimal size of the template in the literature. Such an exhaustive search is responsible for the elevated computational cost of the algorithm, as it increases the runtime proportionally to the number of tried template sizes.^12^ Another factor affecting the computational complexity is the choice of FFT for convolution to compute the NCCC; however, its overall complexity of 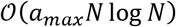 is still costly (*a_max_* is the number of tried template radii), especially when processing large MRI datasets.

In this work, we develop a new computational framework for the NCCC with the 3D Gaussian template, where we approximate the convolution with the Gaussian as multiple convolutions with a box kernel, per the principle of central limit theorem (inspired by a fast method of computing continuous wavelet transform^23^), which follows linear time, 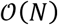, for a fixed radius with respect to the number of voxels in the MRI volume, *N*. This method is fast with the overall complexity of 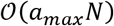, and produces equivalent results compared to the FFT-based algorithm (Eq. (1) – Eq. (15), which have been presented in our preliminary work^12^). We then extend this approach by developing a new template-matching algorithm to compute the NCCC with the optimal-size template in linear time, 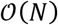, for automatic detection of brain lesions from volumetric MR images. To accomplish this, we propose a mathematical derivation to analytically optimize the maximum template radius for each voxel in the MRI volume, leading to a computational cost that is independent of the radius of template (lesion size). The addition of such a radius-optimized template-matching approach is possible via extending our previous algorithm, but not with the FFT-based method. We tested the performances of our two methods (with overall complexities of 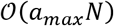 and 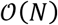) on synthetic data and real MR image databases of MS lesions, and then compared them with those of the conventional FFT-based method and the LPA. Experimental results over synthetic data validate the proposed approach, and lesion-detection results on real MRI volumes show its effectiveness and applicability.

## Methods

### Template Matching in 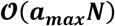

Let 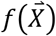 be the *D*-dimensional input image the lesions in which we intend to detect. In this work, the *D*-dimensional Gaussian is proposed as the template, which is approximated – following the central limit theorem – by convolving a *D*-dimensional symmetric and normalized box kernel, *h_D_*(·), with itself *n* − 1 times, denoted as 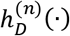 (aka B-splines^24^). In our experiments, a small value of *n* = 2 or *n* = 3 turned out to be sufficient. The sizes of the box kernel and the engulfing template are 2*a* and 2*b*, respectively, in all dimensions; i.e. 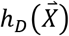 is 1/(2*a*)^*D*^ if 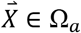, and 0 otherwise, where 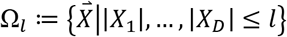. To ensure that the box kernel fits in the engulfing template after the convolutions, we restrict its size as: 0 < *a* ≤ *a*_*max*_ < *b*/*n*. The similarity between the given image and the symmetric template with a varying *a* can be computed from the following formula for the NCCC:

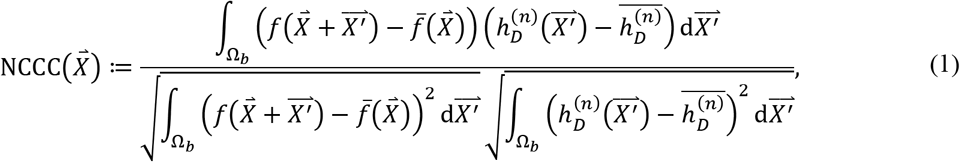

where 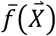 and 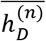 represent the mean of the image inside the template centered at 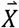, and the mean of the template, respectively. Our goal is to maximize NCCC with respect to both *a* and 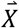, to accurately localize the lesions. In the following, we proceed to calculate different terms in Eq. (1).

Since by definition 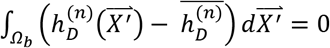, we can omit 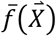 from the numerator of Eq. (1). Noting that 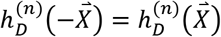, we compute 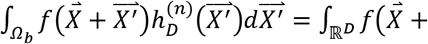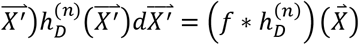. Thanks to the separability property of the template, i.e. 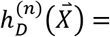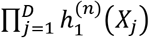, we can first find the solution to this convolution for *D* = 1 and then apply it sequentially for each dimension. For *D* = 1, we note that:

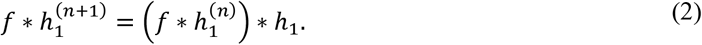

We assume (and then verify) the following solution for the convolution:

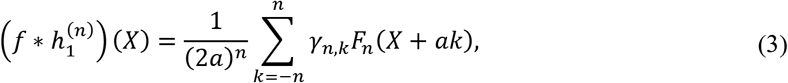

where 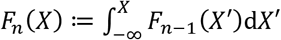, with *F*_0_ ≔ *f*. Let *γ_n,k_* ≔ 0 for |*k*| > *n*, and also *γ_0,0_* ≔ 1 for the case with no convolution. Now, to verify that Eq. (3) is indeed a solution and to compute *γ_n,k_*, we substitute Eq. (3) in Eq. (2):

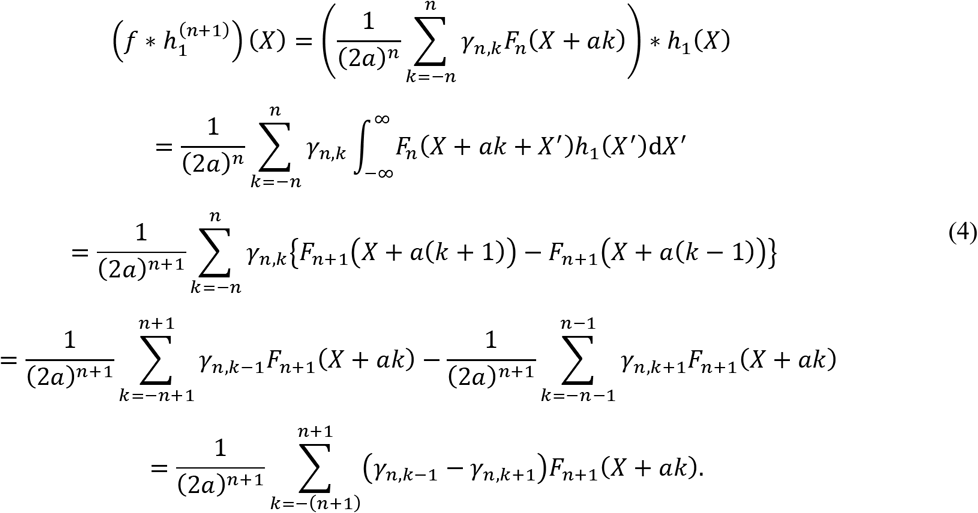

According to Eq. (3), replacing *n* with *n* + 1 should result in:

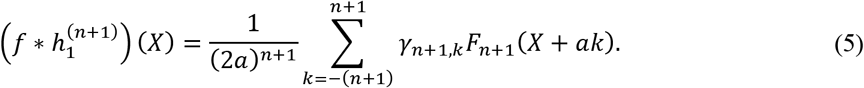

This validates our assumption in Eq. (3) by induction, as Eq. (3) is true for the base case *n* = 0, and provided that it is true for *n*, it holds for *n* + 1 (thanks to similarities between Eq. (4) and Eq. (5)) with the following recursive relationship for *γ*, which is obtained by coefficient matching between Eq. (4) and Eq. (5):

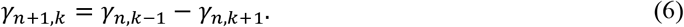

For example, *γ*_1,{−1,0,1}_ = {−1,0,1} and *γ*_2,{−2,…,2}_ = {1, 0, −2, 0, 1}. Being closely related to Pascal’s triangle, Eq. (6) is solved as follows:

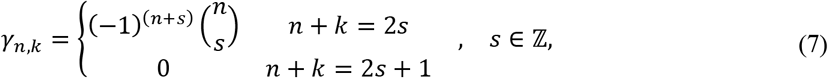

where 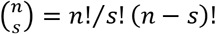 is the binomial coefficient. We now extend this to *D* dimensions and solve the first numerator term of Eq. (1) with the help of *γ_n,k_*:

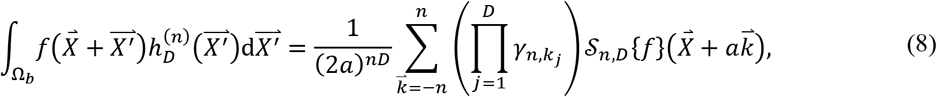

where 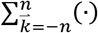 is short for 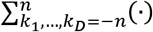. The operator 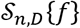 facilitates integration of *f* in all dimensions, and is defined recursively as:

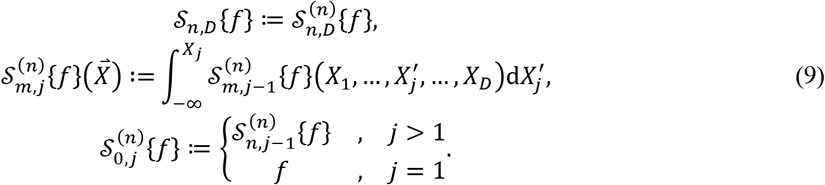

Using the fact that 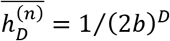 (see below), the remainder of the numerator of Eq. (1) can be computed similarly to Eq. (8) (by fixing *n* = 1):

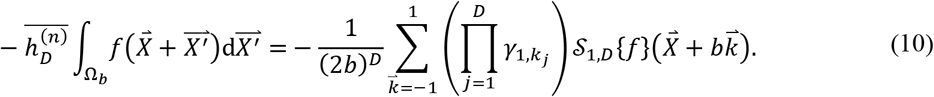

Next, we rewrite the first factor of the denominator of Eq. (1) using the common expansion of the variance as 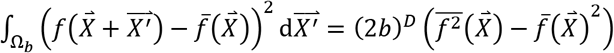, where 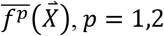, is calculated similarly to Eq. (8) (by fixing *n* = 1) as:

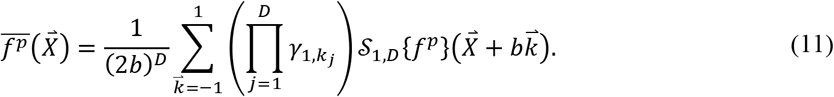

Lastly, we calculate the second factor in the denominator of Eq. (1), again using the common expansion of the variance, as: 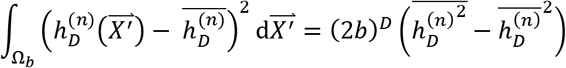. To that end, we calculate the Fourier transform of the box function as 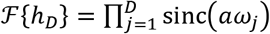, where sinc *θ* = (sin *θ*)/*θ*, which leads to the Fourier transform of the kernel 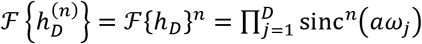 via the convolution theorem. The integral of the template then is 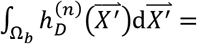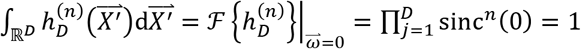, from which the template mean is computed as 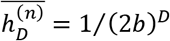. As for the mean of the square of the template, we use Parseval’s theorem as follows:

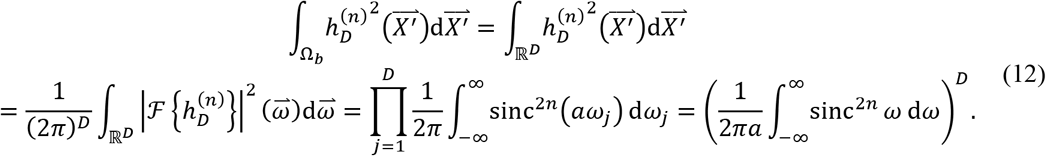

We now define and compute^25,26^:

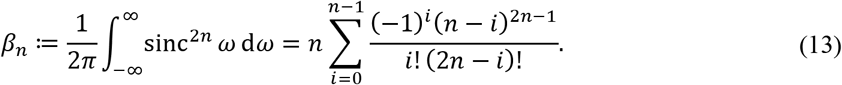

Therefore, 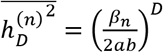, and subtracting 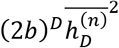 from it leads to:

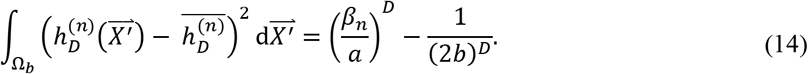

Substituting all the above in Eq. (1) will result in the following formula for the proposed fast approach of NCCC computation:

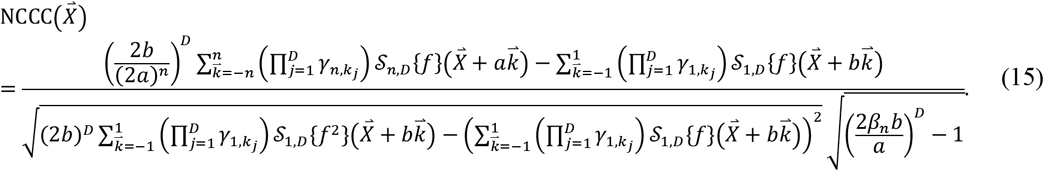

NCCC values range from −1 to 1. In this subsection, we compute the NCCC for several different values of *a*. However, note that 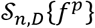 are volumes independent of *a* and can be pre-computed, along with the second term in the numerator and the first factor in the denominator of Eq. (15). Furthermore, the second factor in the denominator is a scalar that is computed fast in 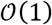 for each *a*. Thus, in the proposed approach to estimate NCCC with the Gaussian template, the bulk of the computational cost is only due to the first term in the numerator of Eq. (15), which can be computed in 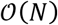 for each *a*, with *N* the number of voxels in the image. On the contrary, the computational complexity of FFT is 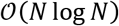 for each *a*,^27^ making the overall computational cost of lesion detection noticeably higher for the FFT-based algorithm (template has to be zero-padded to the size of the image) than the proposed approach. This difference in computational cost is particularly amplified given that the NCCC needs to be repeatedly computed for many values of *a* = 1,2, …, *a_max_*.

### Template Matching in 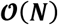

#### Defining NCCC

We intend to optimize the NCCC with respect to the template radius; however, analytical maximization of Eq. (15) with respect to the template radius is difficult. Therefore, we aimed at redefining NCCC, which would be suitable for voxel-wise maximization of template radius. A flowchart of the proposed method is illustrated in Figure 1. Let us assume the template, 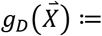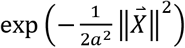, and the ROI are both Gaussian, where ∥·∥ denotes the Euclidean norm. As in the previous section, we solve NCCC by considering all terms of numerator and denominator, individually. We begin by focusing on the first term of the numerator of Eq. (1). For a Gaussian-shaped lesion, this term can be solved as:

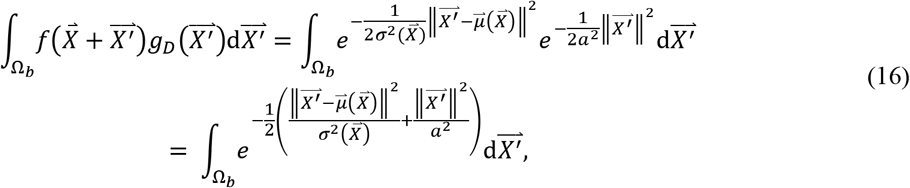

where, 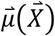 and 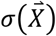 are the mean (with respect to 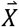) and standard deviation (SD) of the ROI, i.e. the location and the radius of the lesion, respectively. We now simplify Eq. (16) by reducing the power of exponential via completing the square as,

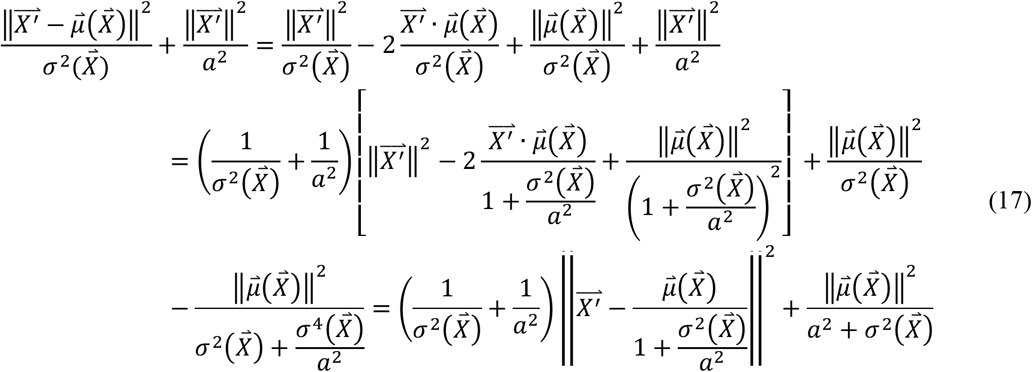

**Figure 1.**
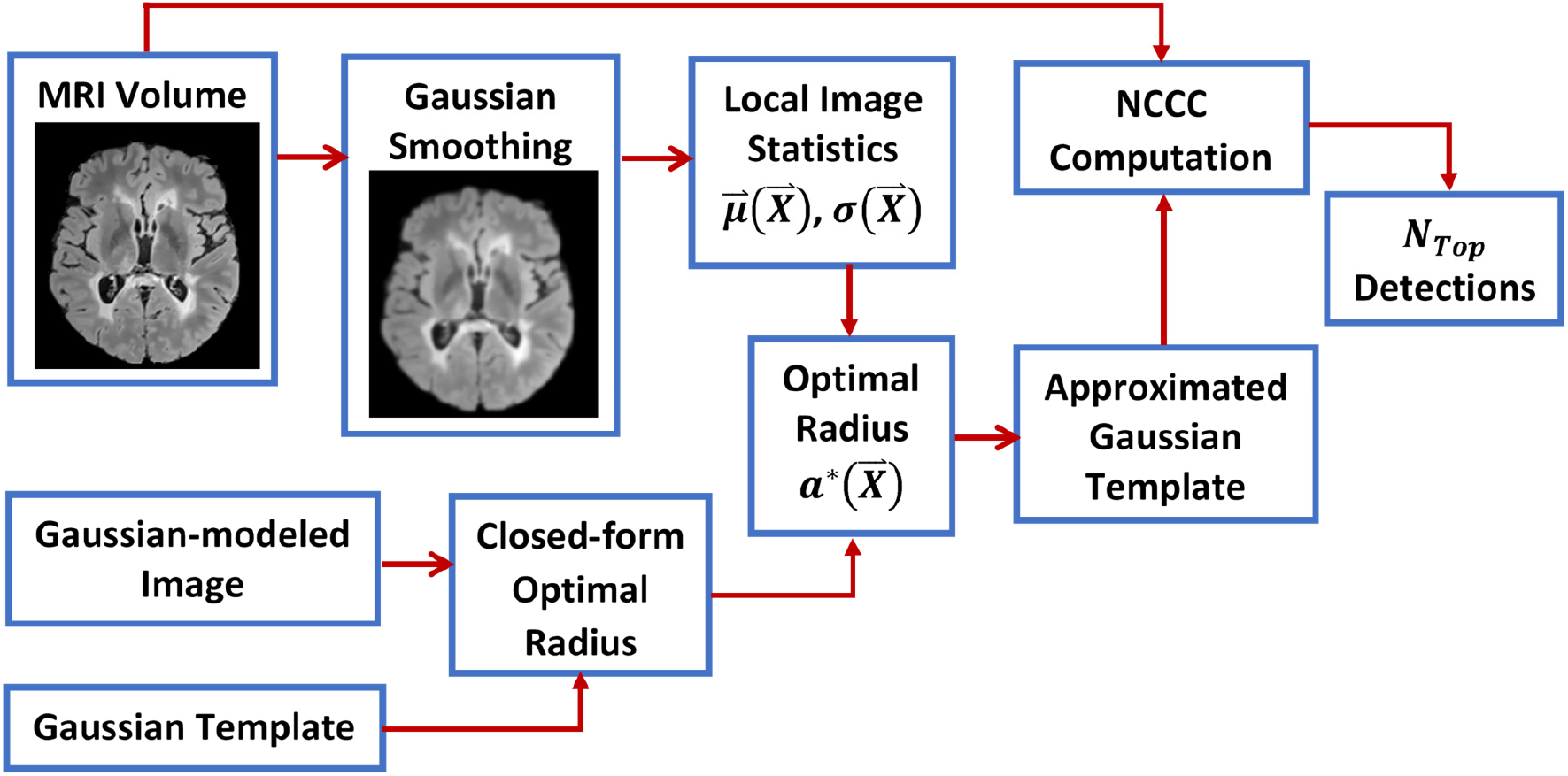
A flowchart depicting the proposed 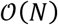 method. We first calculate a closed-form formula for the optimal radius by Gaussian modeling of the lesion, which we then apply to the local image statistics (estimated from a smoothed version of the MRI volume) to compute the optimal radius at each voxel. We then use an approximated Gaussian template (by consecutive convolutions with the boxcar kernel) to compute the NCCC at each voxel, from which we keep a pre-determined number of top detected lesions.

Now we can rewrite Eq. (16) as,

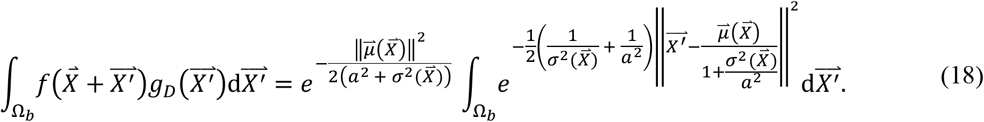

We define 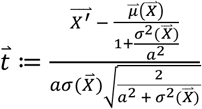; hence, 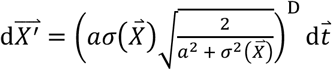. After the change of variables, and assuming the radius of engulfing template to be sufficiently large, i.e. *b* → ∞, Eq. (18) can be reduced by using the properties of the error function 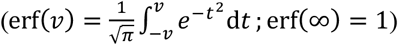 to the following, which completes the calculation of the first term of the numerator:

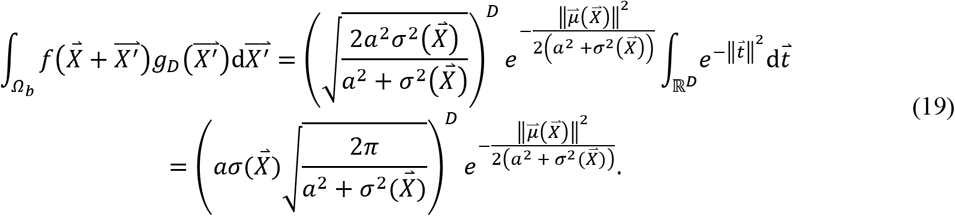

We now focus on the first term of denominator, i.e. image variance inside the template. We use the expansion of variance, 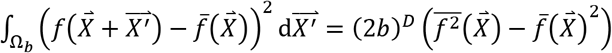, and compute 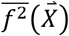 and 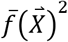 as follows.

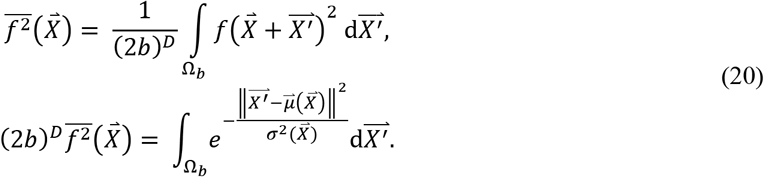

Again, for a large *b* → ∞, Eq. (20) can be simplified with the help of the error function as:

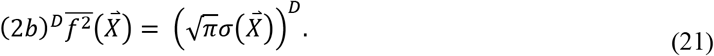

Similarly, 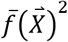 is computed in the same way as in Eq. (20), and for a large *b*, one can see that,

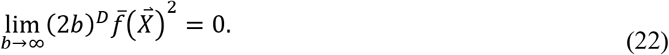

Subtracting Eq. (22) from Eq. (21), we get an approximation for the first term of denominator of NCCC for a large *b* as,

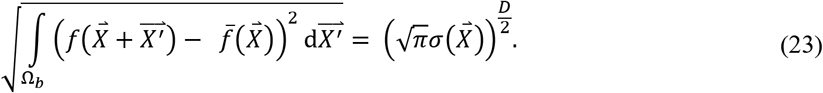

Next, we took on the second term of denominator, i.e. the template variance. We followed the expansion solution to initiate its computation. First, the mean of the square of the template can be computed as,

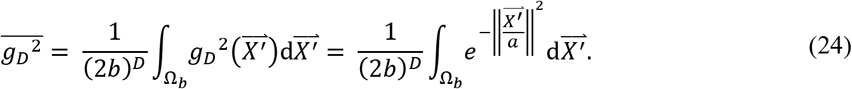

Like previous steps, for a large *b* → ∞, Eq. (24) can be simplified as, 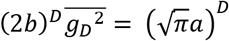. Similarly to above, the mean of the template, 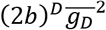, is zero for *b* → ∞, and we obtain the approximate solution (for large *b*) of the second term of denominator of NCCC as,

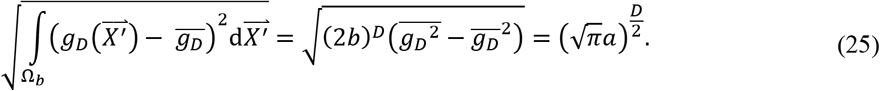

Next, we compute the second term of numerator of NCCC. We have, 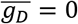; therefore, 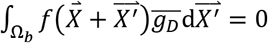.

Substituting the obtained solutions of all terms associated with NCCC in Eq. (1), and simplifying it further, results in the following formula as the final solution for computing NCCC, when both image and template are Gaussian.

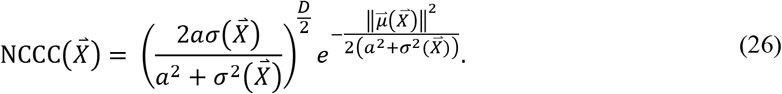

#### Optimization of Template Radius

The present form of NCCC in Eq. (26) is suitable for analytical maximization with respect to the template radius, *a*. The voxel-wise maximizer, *a*, would allow the computation of NCCC in a single step, thus reducing the computational complexity to the linear time, given that the NCCC no longer needs to be computed for a set of template radii. Let us define 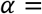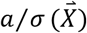. With this notation, Eq. (26) can be simplified as,

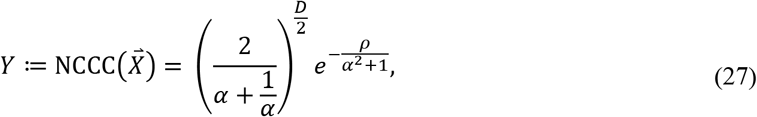

where, 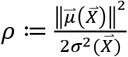. After substitution, we maximize the NCCC by equating the derivative of Eq. (27) with respect to *α*, 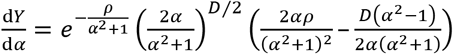, to zero, resulting in the following analytically optimized voxel-wise template radius,

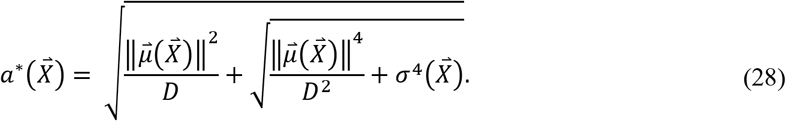

#### Local Statistics of Image Volume

The local statistics, mean 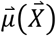 and SD 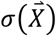 above, are the key parameters to compute the voxel-wise optimized radius. We measured these two parameters from Gaussian smoothed image to counter the small intensity variations in MRI. Gaussian smoothing was performed on the MRI volume with a 3D Gaussian filter of size *W* × *W* × *W*, where *W* = 2*b*′ + 1 and *b*′ < *b*, and an SD of 2 voxels. We then computed the background minimum image, *f_BG_*, via moving minimum filtering over the smoothed image with a kernel of the same size to record the local minimum value inside the sliding window, which we then used in computing local statistics. The time complexity of this step is still kept 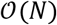 by sequentially applying 1D filters in each dimension in both filtering steps.

##### Mean

We compute the local (moving) mean at each voxel in the smoothed image. The moving minimum is locally subtracted from the local image, in order to remove the effect of the background in the local neighborhood. We define the *D*-dimensional mean as described below.

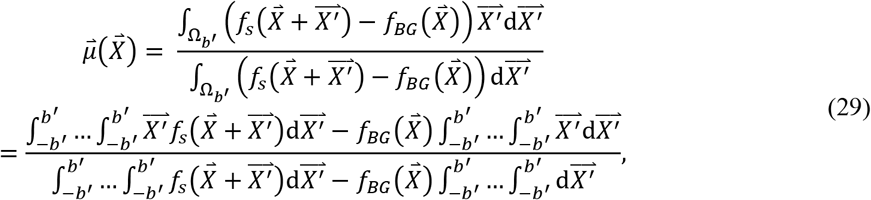

where 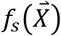 is the Gaussian-smoothed image. The second term of the numerator is zero due to symmetry. To simplify the calculation of the above expression, we make use of a linear change of variables. Assume, 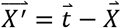, so 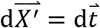. Hence, we can rewrite Eq. (29) as,

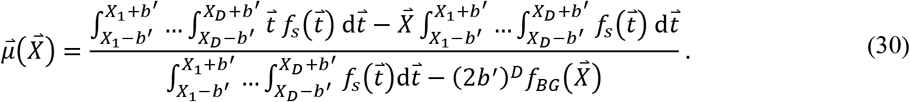

The above equation can be computed in linear time using our proposed solution for convolution in *D* dimensions (refer to Eq. (8)) to solve all the integrals (with *n* = 1) as shown below.

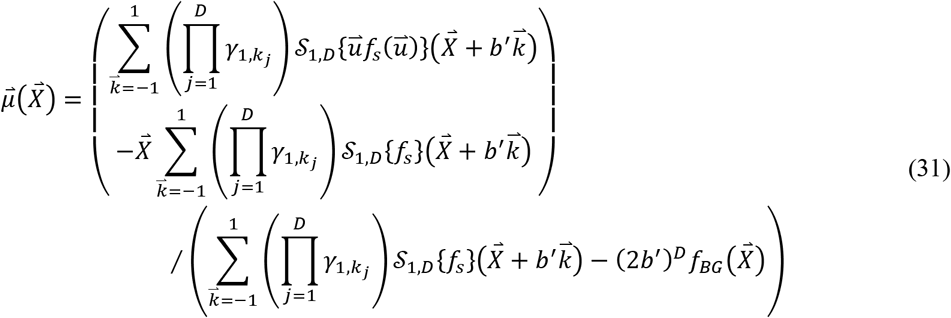

##### Standard Deviation

SD is computed for each voxel by using a *W* × *W* × *W* kernel in Gaussian-smoothed background-subtracted image. From the definition of variance, the SD (*σ*) can be written as,

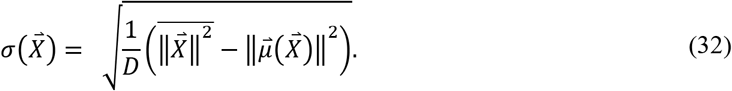

The covariance of a vector is defined as a *D* × *D* matrix. However, given that the template is a radially symmetric Gaussian, we consider a scalar variance here as 1/*D* of the trace of the covariance matrix. We proceed by computing 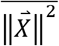 as,

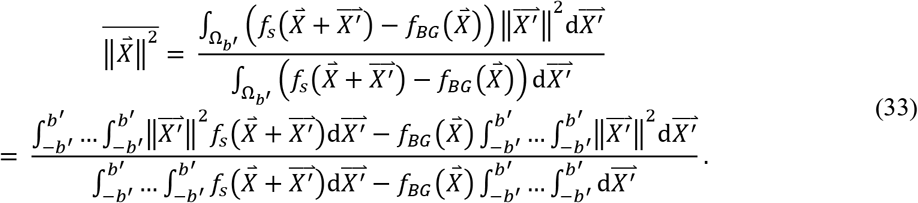

The denominator of 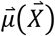 and 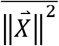 are the same, which is computed in Eq. (31). We compute the first term of numerator of 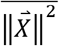, by a change of variables, 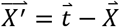, so 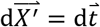,

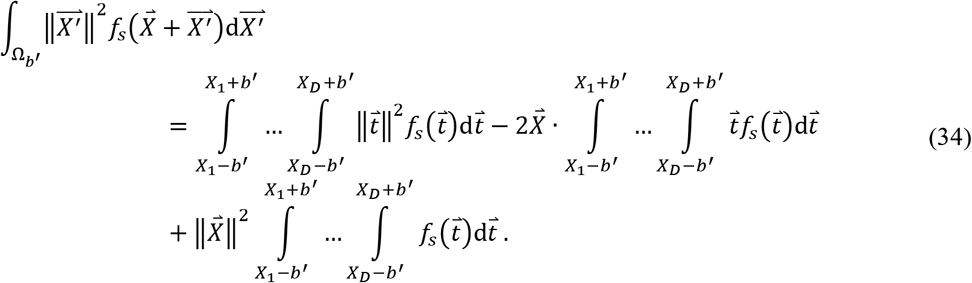

Similarly, to the case of the mean, we apply our approach to computation of convolution in the above equation (with *n* = 1) as described below,

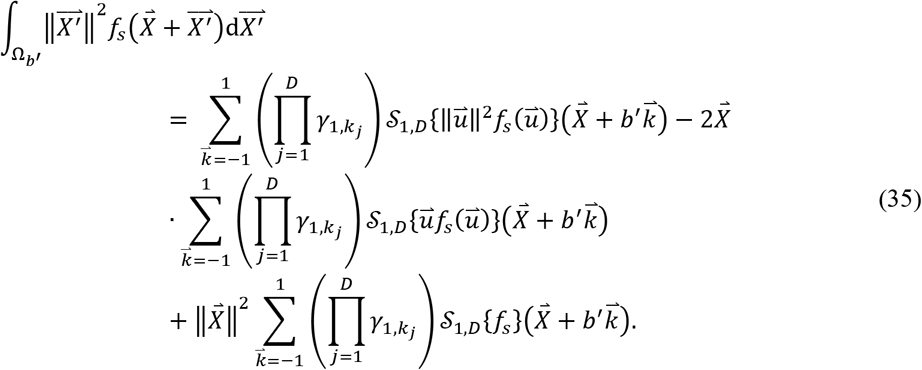

We compute the second term in the numerator of Eq. (33) the same way as in Eq. (35), by replacing 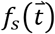 with the constant 1. Finally, we have the solution of the terms necessary to calculate local SD, 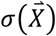, from Eq. (32) in linear time.

#### Template Matching with Maximized Radius

In this step, the mean and SD are substituted in Eq. (28) to obtain the “*a*”-maximized volume. Further, the optimized *a*\* volume should be thresholded as *a*\* ← min(*a*\*, *b/n* − 1) before being substituted in the proposed solution of NCCC as described in Eq. (15). This is how the NCCC computational complexity can be reduced form 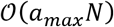 to 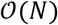, i.e. we no longer need to use a set of templates with varying radii. In contrast, the FFT approach cannot make use of an optimal radius map to reduce its complexity any lower than 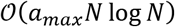, given that FFT-based convolution cannot be performed with a locally varying kernel.

## Results

In this section, we present experimental results comparing the proposed methods and other existing techniques, on 3D synthetic and real MRI data (*D* = 3). We first describe the FFT-based NCCC computation, to which we will compare our methods next.

### FFT-Based Template Matching

We describe here how to use FFT to calculate the NCCC via Eq. (1). We define a truncated Gaussian kernel (n → ∞) as, 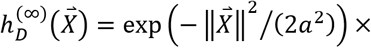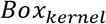 for *a* = 1, …, *a_max_*, which is used as the template. *Box_kernel_* is a volume of equal size to the image, with the value of 1 inside *Ω_b_* and 0 outside. We solve the convolution in the frequency domain by multiplying the FFT of *f* by that of 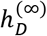 and inverse Fourier transforming (IFFT) it, resulting in the first term of numerator of Eq. (1). The remainder of the numerator is computed similarly by using *Box_kernel_* instead of 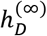. The mean of the template is calculated as, 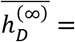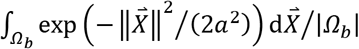. The denominators can be similarly computed. In the FFT-based approach, NCCC needs to be computed for a set of template radii, resulting in the computational complexity of 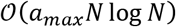.

### Experiment on Synthetic Data

We first evaluated the proposed methods on synthetic data and compared their performances with that of FFT. We compared the detection accuracy and runtime of the three methods. We created twenty artificial volumes of size 513 × 513 × 513 voxels, where each volume contained an enhancing sphere located at a random location with a random radius (varying in between 8 to 21 pixels) to generate the synthetic data. In our first experiment, we aimed to determine how accurately the enhancing spheres could be detected with their approximated radii from the respective volumes. During the experiments, we used the same input parameters for all methods such as *b* = 50 and *a_max_* = 24 (except for the 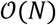 method, instead *b*′ = 24). Note that *n* is ∞ for FFT and 2 for our methods. *a_max_* was calculated as *a_max_* = *b/n* − 1. During this experiment, we observed that both the FFT-based and our 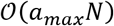 template matching accurately identified the centers of the enhancing spheres in the respective volumes. For the proposed 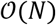 method, we calculated the analytically optimized *a*\* for each voxel, using which we computed the NCCC in linear time over the entire image. We then simply looked for the voxel with maximal NCCC value to identify the center of the detected enhancing sphere, and the *a*\* at that location was considered to be the detected radius. Since the SD of the Gaussian is proportional – but not necessarily equal – to the radius of the detected binary sphere, we conducted a linear regression analysis to reveal the ratio of the gold-standard binary radius to the detected Gaussian SD. From the experiment, we found their ratio to be 1.61 ± 0.05 and 1.74 ± 0.06 for the 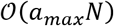 and the FFT methods, respectively, both values significant (*p* = 9 × 10^−18^).^12^

As for the 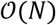 method, we first obtained the abovementioned ratio as 1.96 ± 0.11 (*p* = 5 × 10^−16^). The centers of the enhancing spheres, however, were not detected by the 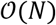 method with pinpoint accuracy; the mean distance between the gold-standard and detected centers was 1.6 ± 1.8 voxels. Consequently, we looked up the values of *a*\* at the known centers of the spheres, and recomputed the gold-standard-to-detected ratio as 2.12 ± 0.11. Next, we repeated the 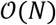 experiment with the corrected *a*\* ← 2.12 × *a*\*. This helped us to reach absolute localization accuracy, with the new gold-standard-to-detected ratio of 1.01 ± 0.03 (p = 8 × 10^−21^).

We performed another experiment on the synthetic data to analyze the runtime. The data were generated by creating 20 different volumes with linearly increasing number of voxels from *N* = 10^6^ (100 × 100 × 100) to *N* = 1.25 × 10^8^ (500 × 500 × 500), each of which containing a homogeneous bright sphere (all ones) of radius 17 pixels at the center. We ran the scripts of the algorithms 10 times for each synthetic volume for sphere detection with the same parameters as above. We measured the mean and SD of the 10 runtimes and compared them among the different methods, as depicted in Figure 2. We ran the experiments in MATLAB on a Linux compute cluster with each node including 8 processors with 56 GB shared virtual memory (cluster allots 1 CPU with 7 GB of virtual memory to a single submitted job). To run the scripts on single thread for an unbiased comparison, we used the `singleCompThread` option, but we found that MATLAB still sometimes multithreaded the codes on multiple cores. Therefore, we report both the wall time (the real-world time elapsed between the start and end of the code) and the CPU time (the total time spent by all cores to execute the script). The 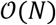 method (green curve in Figure 2) was respectively 3.3 ± 0.7 and 1.4 ± 0.1 times faster than the FFT and 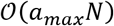 methods in terms of CPU time, and 2.2 ± 0.6 and 1.2 ± 0.2 times faster in terms of wall time. Additionally, the 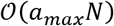 approach was 2.5 ± 0.4 and 1.9 ± 0.4 times faster than the FFT method in terms of CPU and wall times, respectively. The non-monotonicity seen in the FFT runtime is partially because the radix-2 implementation is faster when each dimension is a power of 2. We also observed that the 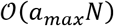 method required 1.7 ± 0.3 and 2.5 ± 0.3 times less resident memory then the FFT and 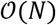 methods, respectively, and that the FFT-based template matching used 1.5 ± 0.3 times less memory than the 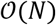 method.

**Figure 2.**
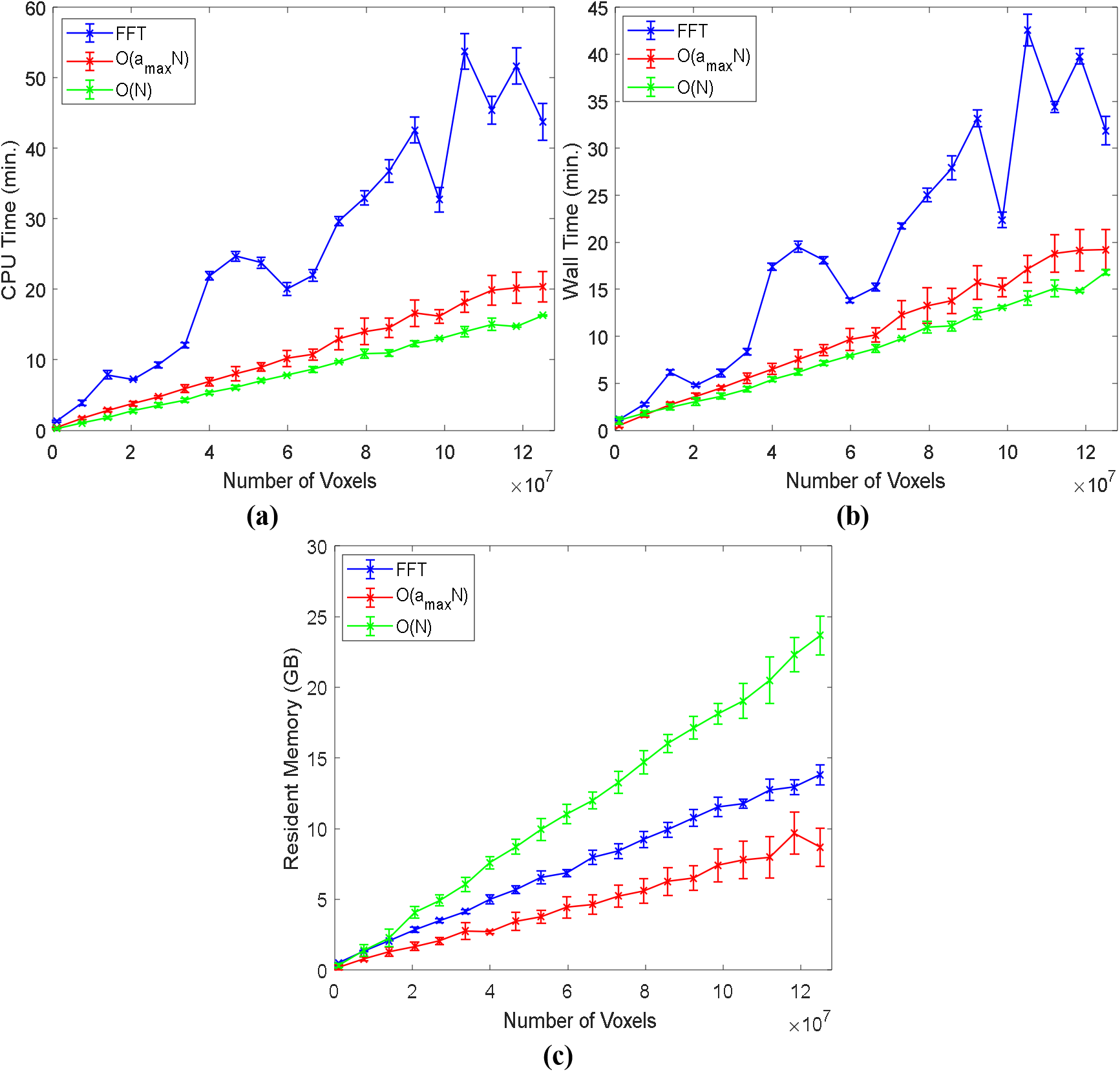
Analysis of runtime (CPU and wall time) and resident memory of 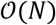 (green), 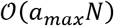 (red), and FFT-based (blue) approaches on synthetic data. As expected, the 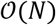 approach is the fastest of the three, with a linear trend with respect to the number of voxels. The runtime of the FFT method fluctuates due to the necessary zero-padding to the nearest power of 2.

### Experiments on Clinical MRI

We tested all three methods on *in vivo* MRI data of human brain for automatic detection of multiple sclerosis (MS) lesions. MS is a chronic demyelinating disease that generally affects the central nervous system, with lesions appearing in various regions of the brain, predominantly the white matter (WM)^4,28^. Given that the cerebrospinal fluid (CSF) is attenuated in FLAIR (though sometime lesions are iso-intense to the hyper-intense gray matter (GM)), FLAIR MRI is suitable for understanding the lesion burden and measuring the amount of damage due to demyelination^3,6^. We tested the methods over the FLAIR sequence of two different MS lesion MRI databases; the first one is the training data of ‘MS lesion segmentation challenge 2008’ (MSLS 2008), which was conducted in conjunction with MICCAI 2008 (http://www.ia.unc.edu/MSseg), and the other dataset is the training data of the ‘MS segmentation challenge using a data management and processing infrastructure’ (MSSEG 2016), which was conducted in MICCAI 2016 (https://portal.fli-iam.irisa.fr/msseg-challenge/overview).

### MRI Scans of MS Subjects

We used 20 cases of training data of MSLS 2008 to evaluate the methods using the gold-standard annotation from the Boston Children’s Hospital (BCH) rater as the reference. Isotropic-voxel (0.5×0.5×0.5 mm^3^) FLAIR images of size 512×512×512^29^ were preprocessed through N4 bias field correction (https://www.slicer.org), brain extraction from T1 images using the volBrain tool (http://volbrain.upv.es), and then the resulting brain masks were multiplied with bias-corrected FLAIR images to remove the skulls.

The 15 cases of preprocessed (nonlocal means (NLM) filtering, rigid registration, skull stripping, and N4 bias field correction) MSSEG 2016 training data, as provided in the challenge website, were used in this study.^6^ We sampled the 3D FLAIR images in the slice direction of each volume (with linear interpolation for FLAIR and nearest-neighbor for the gold-standard lesion and brain masks) to obtain isotropic-voxel (1×1×1 mm^3^) data. After sampling, images were transformed into the new sizes, including 315×512×512, 154×224×256, and 245×336×336. The consensus segmentation obtained from all seven delineations (by seven expert raters) was used as the gold-standard for the performance evaluation of all methods implemented in this study.^6^

We min-max normalized the images of both databases before all experiments, and also kept the same input parameters during the experiments, such as *b* = 18 and *a_max_* = 8 for the 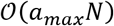 and FFT-based methods, whereas the 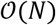 method dealt with all similar parameters, except *a_max_*; instead we had *b*′ = 8. We chose *n* = ∞ for the FFT methods and *n* = 2 for the other two methods, but eventually following a similar approach to find *a_max_* in the FFT and 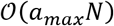 algorithms.

### Performance Evaluation on Clinical MRI Data

The performance of all methods applied for lesion detection were quantified with respect to the expert-made manual annotation of MS lesions in MRI volumes by evaluating the popular statistical metrics such as the true positive fraction (TPF, or sensitivity) and false positive fraction (FPF, or false discovery rate)^9^ as defined below,

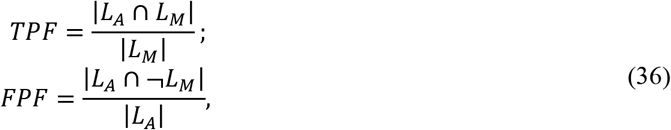

where, *L_A_* and *L_M_* are the label volumes corresponding to automatic detection and manual annotation, respectively, and ¬ denotes negation. A TPF value close to 100% signifies that all voxels representing lesions in the annotated gold-standard are correctly detected, while the 0 value indicates absolutely no detection. Conversely, an FPF value of 100% means no detection, and a value of 0 means that all detected voxels were correctly inside the label.

We now discuss how we used the above metrics to evaluate the performance of the 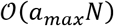, 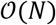, and FFT approaches. We first computed the *a*-maximized NCCC from the FLAIR volume using each method. Many methods in the literature threshold the NCCC to keep putative lesion voxels, on which various performance metrics are evaluated.^10,17,20^ In this work, instead of selecting a threshold, we considered *N_Top_* detected voxels with the highest NCCC value. We found the voxel with maximal NCCC from the resulting *a*-maximized NCCC volume, masked a sphere around it with a radius twice the optimal value of *a*, updated the NCCC volume by filling the sphere with the value of −∞, then repeated this process to find all *N_Top_* maxima.

Next, as described in Algorithm 1, we created a sphere of all ones with the radius 1.61 times the optimally detected *a*, centered at the location of top detection, inside an all-zero volume, *M*, with the same size as the input image. We compared *M* with the gold-standard label to calculate the TPF and FPF for the case of *N_Top_* = 1. We then added an enhancing sphere in *M* for the next detection to compute the TPF and FPF for *N_Top_* = 2 and so on.

**Algorithm 1:**
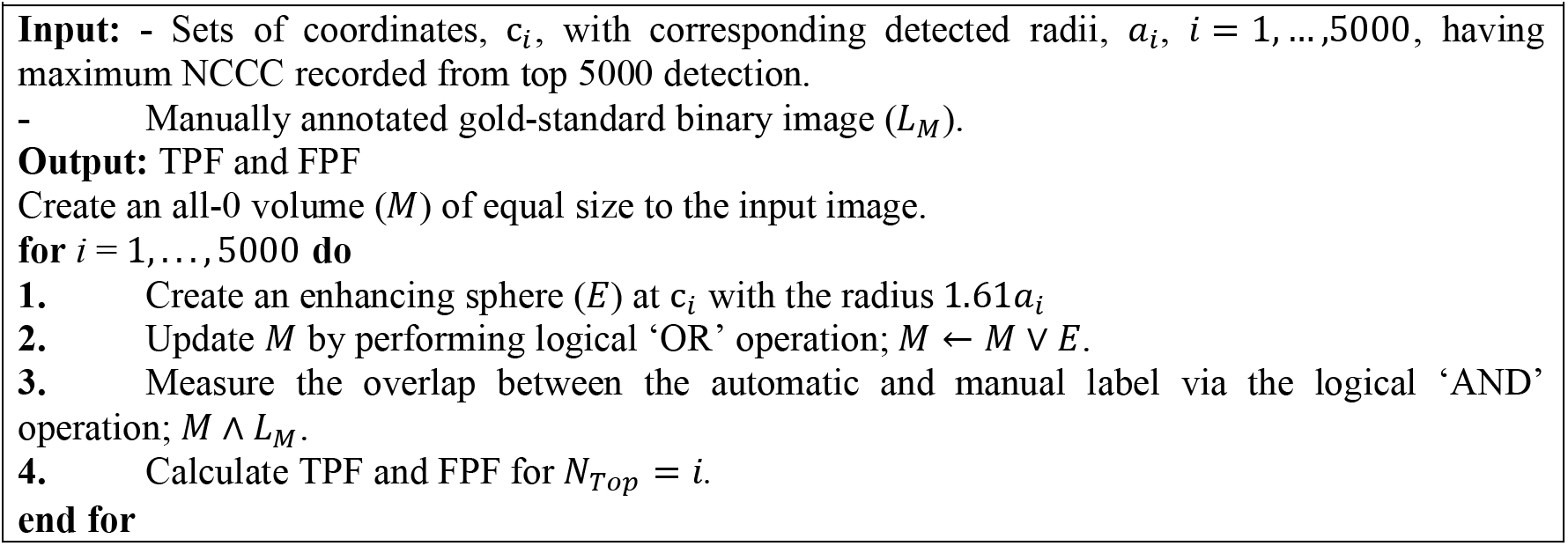
Performance evaluation of algorithms for automatic lesion detection from brain images

The mean TPF vs. FPF graphs over the 20 subjects of MSLS 2008 and the 15 subjects of MSSEG 2016 are presented in Figure 3 for the three methods. We also recorded the computation times and resident memory usage of the three algorithms while running them over the real data as summarized in Table 1. The runtimes and required memory while executing the entire MATLAB scripts (termed as *Algorithm*) and those of the script till getting the *a*-maximized NCCC volume (termed as *NCCC*), are shown in Table 1. All the experiments over real data were performed within the same computational setup as used in the synthetic data analysis. The MATLAB scripts of all three methods were run only once on each image. The visualization of the top 30 detected lesions resulting from all three template matching algorithms are represented in Figures 4 and 5.

**Table 1.** Summary of computational analysis over real MRI database.

| SUMMARY | | | FFT | $\mathcal{O}(a_{max}N)$ | $\mathcal{O}(N)$ |
| --- | --- | --- | --- | --- | --- |
| MSLS 2008 | CPU TIME (MIN.) | ALGORITHM | $194 \pm 52$ | $147 \pm 15$ | $163 \pm 18$ |
| | | NCCC | $16.6 \pm 0.3$ | $8.8 \pm 0.8$ | $17.0 \pm 0.8$ |
| | WALL TIME (MIN.) | ALGORITHM | $195 \pm 52$ | $180 \pm 15$ | $164 \pm 18$ |
| | | NCCC | $23 \pm 3$ | $21 \pm 1$ | $19 \pm 1$ |
| | RESIDENT MEMORY (GB) | ALGORITHM | $17.1 \pm 0.4$ | $12.14 \pm 0.02$ | $24 \pm 1$ |
| | | NCCC | $16.8 \pm 0.8$ | $9.91 \pm 0.03$ | $23.7 \pm 0.9$ |
| MSSEG 2016 | CPU TIME (MIN.) | ALGORITHM | $45 \pm 37$ | $48 \pm 39$ | $46 \pm 38$ |
| | | NCCC | $5 \pm 4$ | $3 \pm 2$ | $5 \pm 4$ |
| | WALL TIME (MIN.) | ALGORITHM | $45 \pm 37$ | $49 \pm 39$ | $47 \pm 38$ |
| | | NCCC | $5 \pm 4$ | $3 \pm 2$ | $8 \pm 5$ |
| | RESIDENT MEMORY (GB) | ALGORITHM | $6 \pm 4$ | $4 \pm 3$ | $7 \pm 5$ |
| | | NCCC | $5 \pm 4$ | $3 \pm 2$ | $7 \pm 6$ |

**Figure 3.**
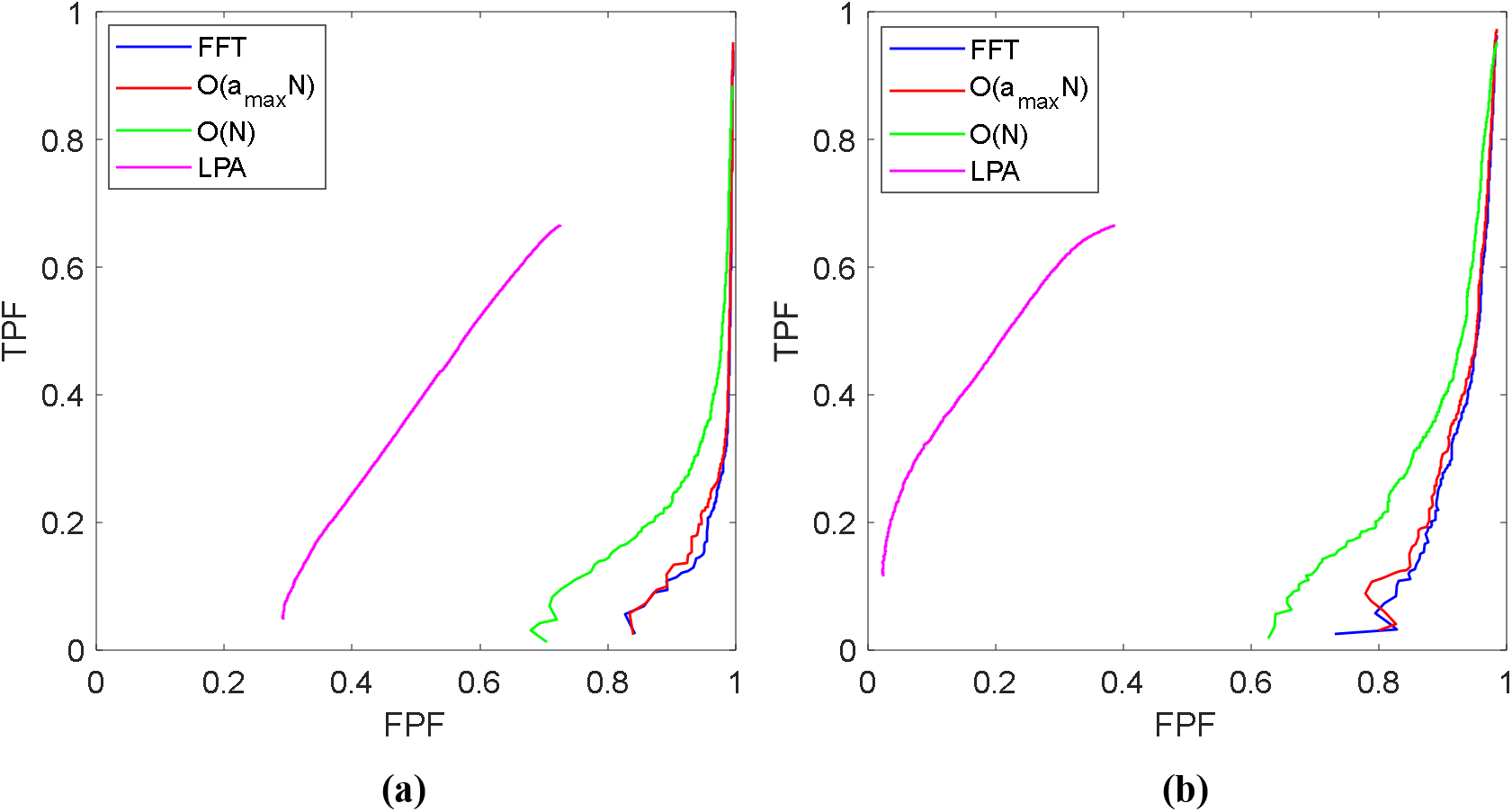
Performance analysis (TPF vs. FPF) of proposed 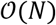 (green), 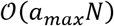 (red), and other FFT-based (blue) and LPA (magenta) approaches over real MRI image database of (a) MSLS 2008 and (b) MSSEG 2016. The 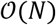 method has the highest accuracy among the three NCCC-based approaches. Note that the superior performance of the LPA approach is expected, given that it is *supervised* segmentation, as opposed to *unsupervised* template-matching lesion detection.

**Figure 4.**
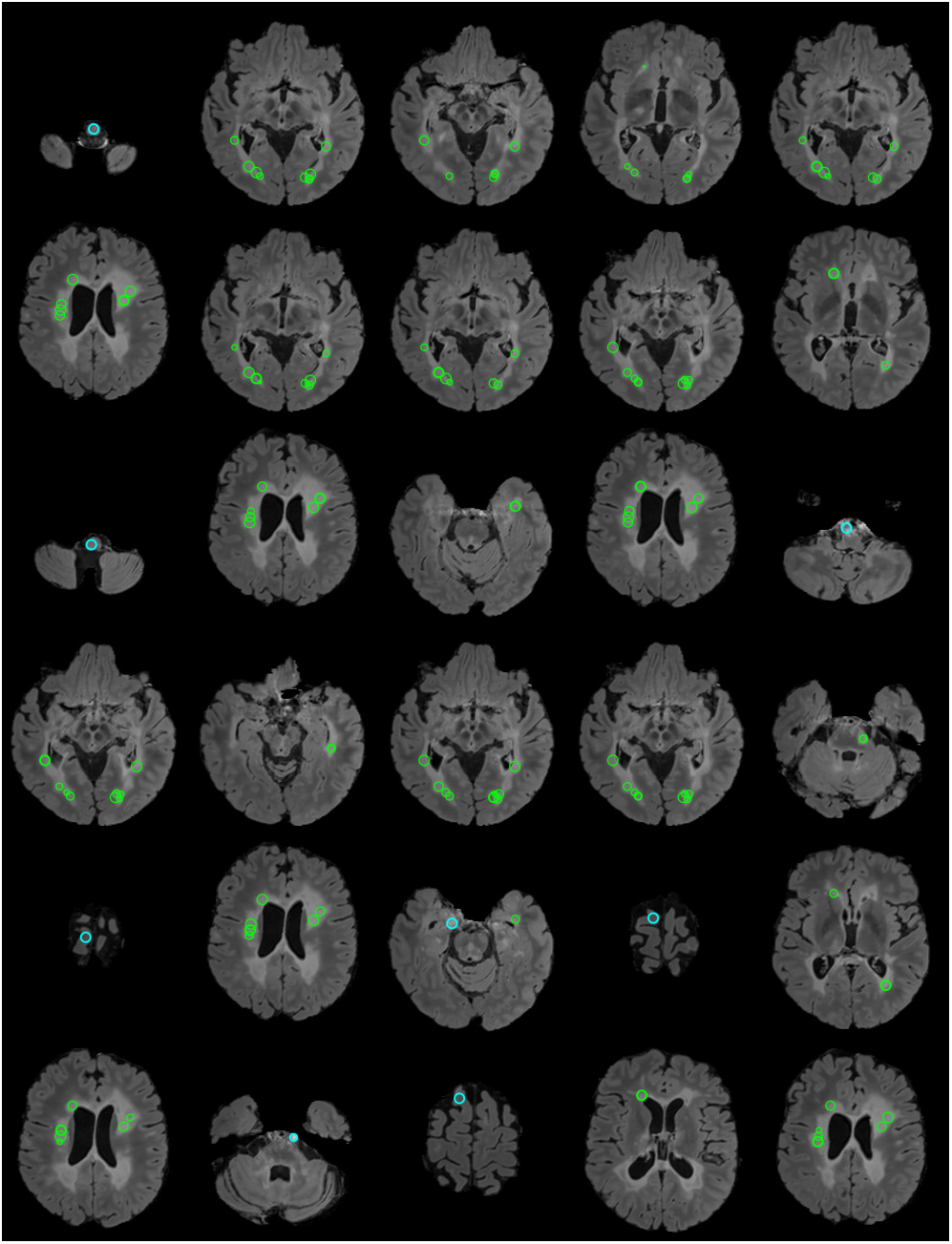
The visualization of the top 30 detected lesions resulting from the proposed 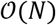-based template matching algorithm on the first case of MSSEG 2016 training data. The green and cyan circles indicate the true and false positives, respectively. Thick and thin circles are detected in the shown and nearby slices, respectively. The top-30 detections are mostly true positives and in the white matter region. False positives are often hyperintense regions mimicking lesions.

**Figure 5.**
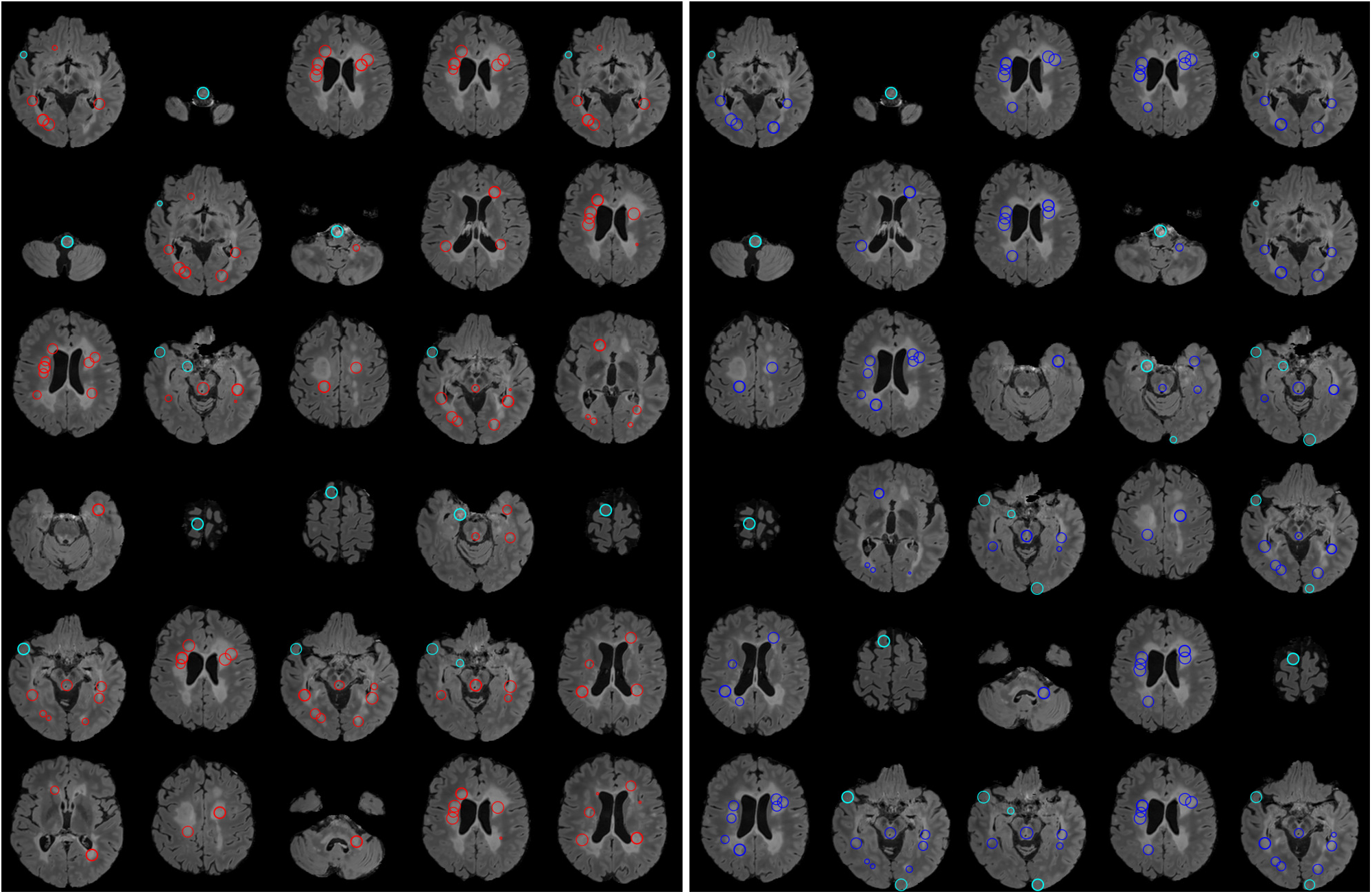
The top 30 detected lesions resulting from the 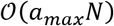- (left) and FFT-based (right) template matching algorithms on the first case of MSSEG 2016 training data. The red 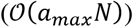 and blue (FFT) circles indicate the true lesions, and the false lesions are identified with cyan colored circles. Thick lesions are those centered at the shown slice, whereas thin circles represent the intersections with other detected spheres in the nearby slices. The performance of the two methods are similar, given that they both exhaustively search for the lesion size that maximizes the same NCCC, albeit computed with different algorithms. However, as seen in Figure 3, these two methods do not achieve the accuracy of the 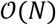 method.

### Comparison with a State-of-the-Art Technique

We compared our methods with the lesion prediction algorithm (LPA)^5^, which is a logistic-regression based supervised learning technique (fundamentally different from our methods) and requires only the FLAIR image with no input parameters for MS lesion detection. We performed binary thresholding over the resulting volume of lesion probability map to compute the TPF and FPF. To maintain the parity with the proposed method in the performance evaluation, we chose the threshold in between 0 to 1 with an interval of 2 × 10^−4^. The LST toolbox (https://www.applied-statistics.de/lst.html) was utilized in statistical parametric mapping (SPM12; https://www.fil.ion.ucl.ac.uk/spm) for the implementation of LPA. The mean TPF vs. FPF plots over the MS lesion databases are illustrated in Figure 3. We can see the better performance of LPA especially on the MSSEG 2016 data compared to the proposed and other methods used in this study.

## Discussion

This paper focuses on fast template-matching based automatic brain lesion detection from MRI volumes. The proposed methods have complexities of 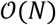 and 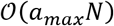, compared to 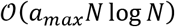 of the FFT approach. From the results of time complexity experiment over synthetic data (see Figure 2(a-b)), it is clear that 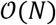 method is the fastest of the three template-matching algorithms for sphere detection. We derived a new analytical solution to obtain the voxel-wise optimal template radius from local image statistics. The proposed analytical solution is a computational alternative to exhaustive search for the template size. The 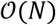 method could detect the bright spheres at their exact centers in the synthetic data (after using the appropriate multiplier). On the other hand, the FFT and 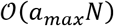 based template matching with exhaustive search possess similar accuracy for detection of the bright spheres as expected. The only small deviation persisting between them for the prediction of the radius of the sphere is due to employment of approximated Gaussian template in the 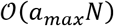 technique. The results on synthetic data validate the theoretical prediction of achieving the linear complexity in template matching by our methods (Figure 2(a-b)). Regarding the resident memory usage, the 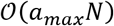 method appears to be quite efficient holding less memory than the other two methods. The 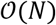 method, however, uses high memory. The analytically optimized template radius can be computed from mean and SD, which are computed from the local 3D neighborhood. The 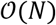 method, therefore, requires storing auxiliary variables (such as mean and variance volumes) in the memory, thereby using more memory than the other two methods. In the exhaustive-search cases, we performed *loop* operation inside the MATLAB script to execute the similarity measure computation step for a set of template radii. We reduced the memory requirement by updating NCCC with the voxel-wise maximized values, thereby avoiding the storage of a separate NCCC volume for each tried radius. Figure 2(c) shows that that the memory requirement increases linearly with the number of voxels in the volumes. In case of real data, the results follow a similar trend. The FFT method requires more memory than the 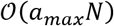 method does, because during template matching with the set of varying radii (*loop* operation) in the FFT method, we need to create a convolution kernel (template) that is as large as the input image, for every tried radius (so they can be multiplied elementwise in the frequency domain).

Regarding the runtimes of the methods on real MR images, Table 1 shows that the CPU time of the 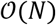 algorithm is higher than that of the 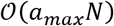 method, and close (till NCCC computation) to that of the FFT-based template matching on the MSLS 2008 data, and that 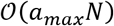 is the fastest of the three on the MSLS 2008 data. These results, which contradict those on the synthetic data, are because NCCC was computed for a set of only 8 different radii for both the 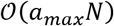 and FFT methods and this *loop*-based operation was proven to be faster than the computation of 10 more convolutions for optimizing the template radius in the 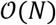 algorithm. In contrast, for a large set of template radii as considered in the experiments on synthetic data, the 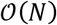 method outperformed the others. The CPU times of the MSSEG 2016 experiments were very close for the three algorithms, since for small images, the gap between *N* log *N* and *N* becomes narrow. Overall, the 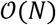 method is expected to run faster for large MRI volumes, or for the detection of larger lesions. *Ex vivo* imaging of human brain, which has attracted researchers to explore microstructural neuroanatomy, can produce brain images with (0.1mm)^3^ isometric voxel resolution.^30^ As the resolution increases, the ROI volume size increases proportionally, necessitating the reduction in computational complexity of the algorithms for processing very large MRI volumes to automatically detect lesions. Note that in real clinical scenario, we do not need to test the detection for top 5000 lesions, instead it would be for fewer numbers depending on the characteristics of the lesions and the clinician’s needs. Also note that the computation of the *a*-maximized NCCC can be seen to take up only a small fraction of the entire runtime (see Table 1), and has more variability.

Prior studies on MS lesions, as discussed in the next section, however, generally segment the lesions and report the detection accuracy from their automatic delineation. Therefore, our approach of lesion detection is different from most available tools. During the performance evaluation, we observed the monotonically increasing characteristics of both TPF and FPF with *N_Top_* for all three methods (except LPA). Thus, a high sensitivity can be achieved with large *N_Top_*, which can be interpreted quite easily from Figure 3. The mean TPF vs. FPF plots (Figure 3) show very similar performance by FFT and our 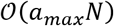 methods over both MRI databases, similar to synthetic data. We can also infer from Figure 3 that the performance graph of the 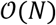 method is slightly better than FFT and 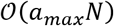, making this method a coherent assistive tool. In addition, the practicality of linear-time template matching can be well appreciated in processing of large 3D volumes with a high *b* (e.g., *b* = 50 showed the efficiency of 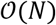 in synthetic data). Throughout the experiment with the 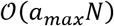 and 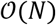 methods, we used an approximated 3D Gaussian template, which is almost radially symmetric, and expected to work robustly with spherical or close-to-spherical lesions. On the contrary, the MS lesions are arbitrary in shape and size (largely varying).^31^ Therefore, the algorithms might not detect the exact centers of such arbitrarily shaped lesions in FLAIR images. This reduced the overlap between the detected region and the gold standard, even when the lesions were correctly localized. This may be one of the reasons behind the rise of false positives. In the context of having large false positives, we would like to mention the work of Cabezas et al.^9^. Their unsupervised methodology deeply depends on tissue priors to estimate the mean and SD of the GM distribution in FLAIR images, for defining a threshold in order to segment the MS lesions. They reported a mean FPF higher than 90% (with close to 40% of mean TPF), as accuracy of lesion detection, without using any false detection refinement step. So, the presence of high FPF seems to be common among the performance of the algorithms, especially targeted to lesion detection from MS subjects, which we will discuss more in next section. Contrary to the unsupervised approaches, the prior knowledge learnt by LPA during the training phase seems to have been effective in improving the performance of this supervised-learning method. This analysis helps in tallying the performance of supervised and unsupervised techniques (Figure 3). In template-matching techniques, a general trend is to fix a threshold over the spectrum of NCCC values so as to evaluate the performance. However, the optimal NCCC threshold may depend on acquisition parameters, and fixing it might lead to the detection performance strongly depending on the threshold. In contrast, our strategy of using maximum NCCC from *N_Top_* detection is not threshold-dependent.

### Visualization of the Detected Lesions

We display the results for top 30 out of 5000 detections in Figures 4 and 5. Blue, red, and green circles represent the FFT-, 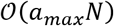-, and 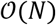-based methods, respectively, on the 2D axial slices of the FLAIR volumes. We present the slices in order of lesion detection. (There may be slice repetition due to multiple detections in the same slice.) Thick circles indicate that the centers of the detected spheres are located in the shown slice. Thin circles represent the intersections between the shown slice and the spheres representing top-30 detected lesions centered in nearby slices. Therefore, the lack of a circle around a lesion means that it has not been in the top 30 (but the algorithm may still have detected it). A limitation that we observed for all techniques is the multiple detection of parts of a large lesion, such as those scattered around the lateral ventricles or on the corpus callosum, which are common in MS^31^. When lesions of various sizes and shapes appeared mainly in the WM^31^, multiple lesions were detected over the same large lesion because of the small *b* that we used. This issue may not be expected to arise in metastatic-lesion detection. The selection of a large *b*, on the other hand, may cause the algorithm to miss smaller lesions. We can observe the presence of false lesions (cyan circles in Figures 4 and 5) included in the top 30 detection for all methods. False positives are mostly in the GM, which is hyper-intense in FLAIR images and sometimes confused with lesion by the algorithms. This is the main cause of high FPF in template-matching algorithms. We also found false positives in other brain regions such as the splenium of corpus callosum, hippocampus, and medulla oblongata. The estimated lesion radius in the 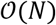 method is comparatively less than FFT- and 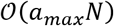-based methods, which is expected from the gold-standard-to-detected ratio radius (1.96 ± 0.11) obtained on synthetic data experiment.

### Comparison with Existing Literature

In this section, we first compare the performance among the existing state-of-the-art methods dedicated for MS lesion segmentation with MSLS 2008 data. We have considered some of the well performing techniques of the last decade (summarized in Tables 2 and 3). Challenge organizers evaluated true positive rate (TPR) and false positive rate (FPR) (with the same definition as TPF and FPF used here) to determine the lesion detection ability^29^ on the test data. Note that the FPR is defined differently here from its standard definition^32^. The lesion voxels are only a small fraction of the MRI volume, thus the standard formula makes the FPR negligible (impactful for small *N_Top_*) due to uninformatively high true negatives. In this study, our method was evaluated on the MSLS 2008 training data with respect to the BCH gold-standards only (performance of existing methods are presented in Table 3 against the BCH gold-standards) due to the unavailability of UNC (University of North Carolina, NC, USA) delineations for all 20 cases. The positive predictive value (PPV) signifies the ratio of the number of true detections to all detections^33^.

**Table 2.**
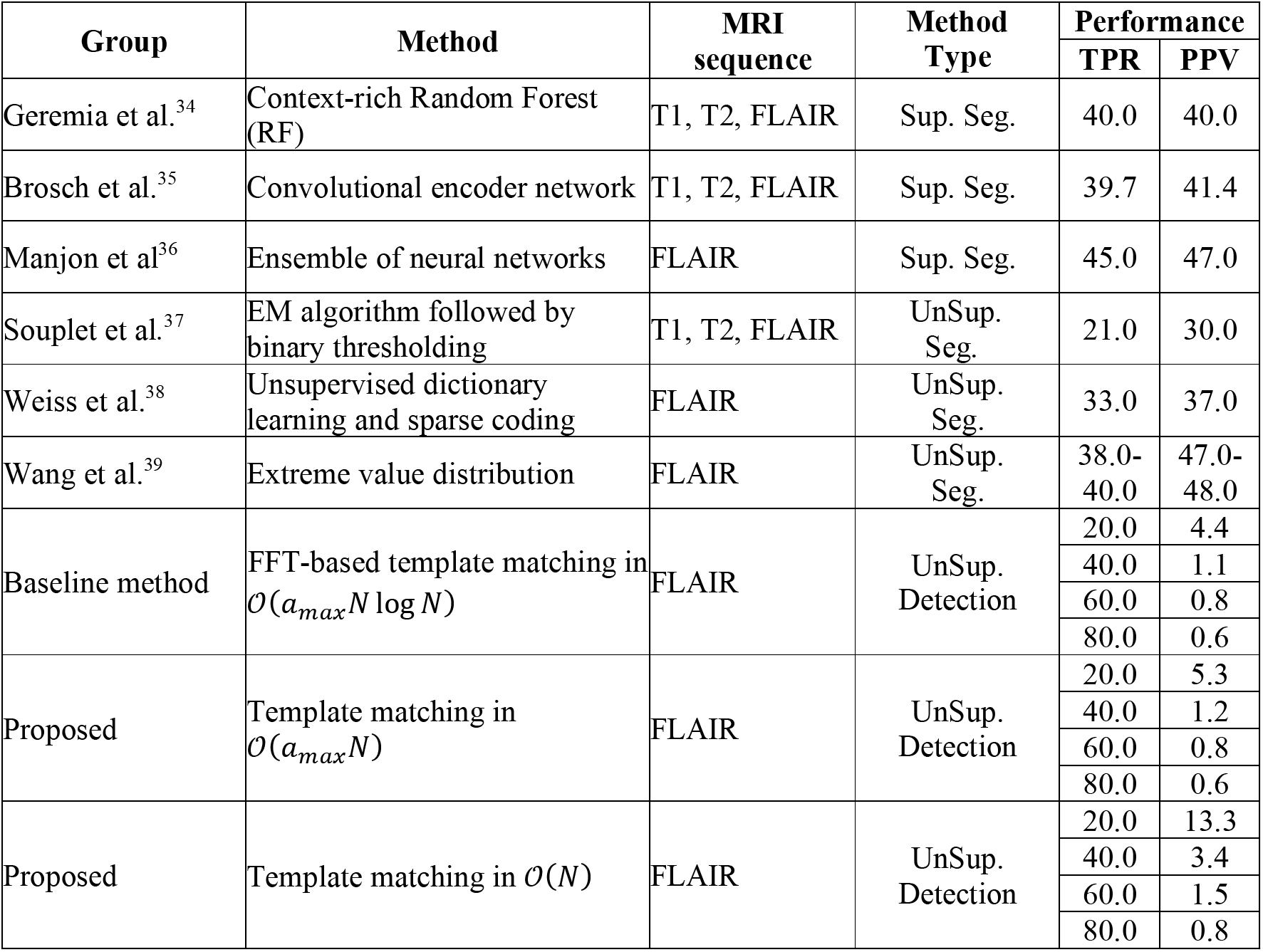
Comparison of the existing Methods on MICCAI MSLS 2008 Training Data^36^. (TPR = TPF; PPV = 1 – FPF.)

**Table 3.** Comparison of Methods on MICCAI MSLS 2008 Test Data^44,51^

| Group | Method | MRI sequence | Method Type | Performance |  |
| --- | --- | --- | --- | --- | --- |
|  |  |  |  | TPR | FPR |
| Geremia et al.<sup>47</sup> | Context-rich RF | T1, T2, FLAIR | Sup. Seg. | 58.0 | 70.0 |
| Guizard et al.<sup>48</sup> | Rotation-invariant NLM | T1, T2, FLAIR | Sup. Seg. | 52.7 | 42.0 |
| Jesson & Arbel<sup>43</sup> | MRF and regional RF | T1, T2, FLAIR | Sup. Seg. | 53.5 | 24.2 |
| Brosch et al.<sup>49</sup> | Deep 3D convolutional encoder | T1, T2, FLAIR | Sup. Seg. | 56.0 | 49.8 |
| Strumia et al.<sup>50</sup> | Geometric brain model | T1, FLAIR | Sup. Seg. | 42.9 | 30.5 |
| Valverde et al.<sup>44</sup> | Cascaded 3D CNN | T1, T2, FLAIR | Sup. Seg. | 68.7 | 46.0 |
|  |  | T1, FLAIR |  | 60.2 | 41.8 |
| Manjon et al.<sup>36</sup> | Ensemble of neural networks | FLAIR | Sup. Seg. | 68.0 | 68.6 |
| Zhan et al.<sup>51</sup> | Multinomial logistic regression with MRF | T1, T2, FLAIR | Sup. Seg. | 44.8 | 56.7 |
| Anbeek et al.<sup>52</sup> | K-Nearest neighbor | T1, T2, FLAIR | Sup. Seg. | 59.1 | 78.5 |
| Jerman et al.<sup>42</sup> | Unsupervised mixture model and random decision forest | T1, T2, FLAIR | Both Sup. and UnSup. Seg. | 71.3 | 62.8 |
| Sudre et al.<sup>53</sup> | Model selection via hierarchical GMM | T1, FLAIR | UnSup. Seg. | 24.6 | 18.2 |
| Tomas-Fernandez and Warfield<sup>54</sup> | Local and global GMM tissue model | T1, T2, FLAIR | UnSup. Seg. | 51.8 | 45.1 |
| Roura et al.<sup>46</sup> | SPM8 and FLAIR thresholding | T1, FLAIR | UnSup. Seg. | 55.4 | 40.5 |
| Ghribi et al.<sup>45</sup> | Atlas-based GMM and lesion expansion | T1, FLAIR | UnSup. Seg. | 84.0 | 69.7 |
| Souplet et al.<sup>37</sup> | EM algorithm followed by binary thresholding | T1, T2, FLAIR | UnSup. Seg. | 57.4 | 68.9 |
| Bricq et al.<sup>55</sup> | Hidden Markov chain | T1, T2, FLAIR | UnSup. Seg. | 46.7 | 51.0 |

In this study, we intended to develop an algorithm for fast and automatic detection of brain lesions, as opposed to determining lesion contours (but still estimating the radius of the detected lesions). In most of the existing works, multi-contrast MR images were considered to delineate lesions^6^, with the exception of some methods that used only FLAIR images (as we did). Using multiple MRI sequences is attractive as it provides a larger feature space than using a single sequence, therefore multi-channel MRIs are widely used in machine leaning and deep neural networks. In this context, our template matching algorithms, focusing on estimating lesion center with an approximated radius, are fundamentally different from the segmentation approaches. During literature review, we observed that Weiss et al. and Manjon et al. avoided computing the whole image to possibly reduce the computational burden, instead they downsampled the image^38^ or used candidate ROIs^36^. We found that MS lesions were detected as regions with outlier intensity values compared to the healthy brain tissue in many works^40,41^. Some research groups designed two-stage segmentation tasks; they first determine the putative lesion voxels with the help of various techniques such as unsupervised mixture models^42^, estimation of lesion and tissue label using Markov random field (MRF)^43^, cascaded convolutional neural network (CNN)^44^, etc., and then segment the lesions by classifying the voxels. In another two-step approach, brain tissues, mainly WM, GM, and CSF, were segmented using different techniques such as atlas-based EM^37^, Gaussian mixture model (GMM)^45^, SPM8/12 segmentation algorithm^46^, etc., before delineating the lesion while making use of the segmented tissues. These multistage approaches generally require multi-contrast MRI. Our proposed techniques compute the image at the original resolution and no spatial prior information on the tissue or lesion or candidate ROIs needs to be extracted; instead, it can directly localize the lesion region based on the NCCC measures, making it a single-stage method. Another key point we observed in the existing literature is the inclusion of post-processing operations, which is a common practice to refine the results and reduce false positives. The presence of flow artifacts in the FLAIR sequence creates hyperintensity in the CSF space, thus sometimes mimicking the lesion^53,56^. In addition, the hyperintense border voxels at the GM-WM junction may result in some false positives^53,57^. As a remedy, various post-processing operators have been adopted, such as morphological operations^37^, setting a minimum lesion volume^46,53,58^, connectivity rules, neighborhood similarity^46^, etc., to reduce false detections and improve FPR and PPV. In this study, we did not perform any post-processing, and only evaluated the raw outcome of the proposed algorithms. The performance of various supervised^34–36^ (Sup. Seg.) and unsupervised^37–39^ (UnSup. Seg.) segmentation techniques, evaluated on MSLS 2008 training data, are shown in Table 2. We also report the results of the proposed methods and the FFT baseline approach for four different values of TPF. One can see that 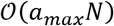 method’s performance is similar to the FFT approach, consistent with the synthetic-data experiments. The 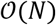 method seems to perform slightly better than the other two template matching methods. But all three methods yield poor PPV (due to high FPF). Without the post-processing step for MS lesion detection, unsupervised detection has previously shown to result in high FPF^9^. Besides the fact that a single image (as opposed to three varying-contrast images) was used here, overlapping lesion intensity, flow artifacts, and typical shape and size of MS lesions are likely causes of high FPF, when lesions are detected from FLAIR images without refinement strategies. Having no user tunable learning parameters, as opposed to (supervised or unsupervised) segmentation approaches, the tested unsupervised detection approaches are not meant to be directly compared to the segmentation approaches presented in Tables 2 and 4 (discussed below), especially given that a detection method is designed to locate – but not delineate – a lesion. Nonetheless, to paint a broad picture of the performances and applicability of existing works, we present all methods in the tables side by side. We also observe from Table 2 that both TPR and PPV are low for all methods, which emphasizes the presence of high false positive with poor lesion detection ability. Table 3 presents a review of the performance of existing supervised^36,42–44,47–52^ (Sup. Seg.) and unsupervised^37,45,46,53–55^ (UnSup. Seg.) segmentation techniques on the test dataset. The values presented in Tables 2 and 3 were retrieved from the literature and leaderboard results (http://www.ia.unc.edu/MSseg/re-sults_table.php), and show that high FPR is very common among the methods even when the computation is done with multi-contrast MRI. Ghribi et al.^45^ pointed out the poor quality of MR images in MSLS 2008 as the reason behind the high FPR of their method. As expected, supervised learning methods performed well in both tables compared to unsupervised techniques.

**Table 4.** Comparison of the existing state-of-the-arts on MICCAI MSSEG 2016 Training Data.^68^ (TPR = TPF; PPV = 1 – FPF.)

| Group | Method | MRI sequence | Method Type | Performance |  |  |
| --- | --- | --- | --- | --- | --- | --- |
|  |  |  |  | DICE | TPR | PPV |
| Mahbod et al.<sup>59</sup> | Multilayer perceptron with morphology-based filtering | 3D FLAIR | Sup. Seg. | 10.2-84.0 | - | - |
| Vera-Olmos et al.<sup>60</sup> | RF classifier and MRF based post-processing | T1, FLAIR | Sup. Seg. | 63.8 | 68.3 | - |
| Salehi et al.<sup>61</sup> | 3D Fully convolutional network with Tversky loss | T1, T2, FLAIR | Sup. Seg. | 56.4 | 56.8 | - |
| Hashemi et al.<sup>62</sup> | Patch-wise 3D fully convolutional DenseNet architecture | T1, T1-GADO, FLAIR, PD, T2 | Sup. Seg. | 69.9 | 78.5 | - |
| Coupe et al.<sup>63</sup> | Rotationally-invariant NLM and patch-wise NLM denoising filter | T1, FLAIR | Sup. Seg. | 72.5 | - | - |
| Chen et al.<sup>64</sup> | Hybrid feature network based on DenseNet architecture | T1, T2, FLAIR | Sup. Seg. | 66.5 | 61.3 | - |
| Kamraoui et al.<sup>65</sup> | DeepLesionBrain (DLB) | T1, FLAIR | Sup. Seg. | 63.9 | 60.8 | 76.8 |
|  | DLB with hierarchical specialization learning | T1, FLAIR |  | 66.9 | 67.1 | 72.8 |
| Valverde et al.<sup>66</sup> | 3D cascaded CNN with $11 \times 11 \times 11$ patch | T1, FLAIR | Sup. Seg. | 44.2 | 42.3 | 61.4 |
| Zhang et al.<sup>67</sup> | 2.5D densely connected fully convolutional network | T1, FLAIR | Sup. Seg. | 66.4 | 65.8 | 74.1 |
| McKinley et al.<sup>68</sup> | 3D-2D CNN (DeepSCAN) architecture | T1, T2, FLAIR | Sup. Seg. | 75.7 | - | - |
| Valverde et al.<sup>44</sup> | Cascaded 3D CNN | T1, T2, FLAIR | Sup. Seg. | 58.7 | - | - |
| Isensee et al.<sup>69</sup> | 2D, 3D, cascade of two 3D U-Net | T1, T2, FLAIR | Sup. Seg. | 74.5 | - | - |
| Beaumont et al.<sup>58</sup> | Multimodal graph cut, EM, and post-processing | T1, T2, FLAIR | UnSup. Seg. | 57.0 | - | - |
| Beaumont et al.<sup>70</sup> | Voxel-wise comparison (GMM and EM) and post-processing | FLAIR | UnSup. Seg. | 50.5 | - | - |
| | | T2 ∪ FLAIR | | 42.4 | - | - |
| | | T2 ∩ FLAIR | | 43.9 | - | - |
| | | FLAIR and T2 ∩ FLAIR | | 56.6 | - | - |
| Beaumont et al.<sup>70</sup> | Voxel-wise comparison without post-processing | FLAIR | UnSup. Seg. | 42.3 | - | - |
| | | T2 ∪ FLAIR | | 29.7 | - | - |
| | | T2 ∩ FLAIR | | 24.7 | - | - |
| Knight & Khademi<sup>71</sup> | Fuzzy classification, thresholding, post-processing | FLAIR | UnSup. Seg. | 60.0 | 53.0 | 80.0 |
| Baseline method | FFT-based template matching in $\mathcal{O}(a_{max}N \log N)$ | FLAIR | UnSup. Detection | 11.0 | 20.0 | 11.8 |
|  |  |  |  | 8.8 | 40.0 | 6.1 |
|  |  |  |  | 6.3 | 60.0 | 3.8 |
|  |  |  |  | 4.4 | 80.0 | 2.4 |
| Proposed | Template matching in $\mathcal{O}(a_{max}N)$ | FLAIR | UnSup. Detection | 11.2 | 20.0 | 12.2 |
|  |  |  |  | 9.5 | 40.0 | 6.8 |
|  |  |  |  | 6.7 | 60.0 | 4.0 |
|  |  |  |  | 4.6 | 80.0 | 2.5 |
| Proposed | Template matching in $\mathcal{O}(N)$ | FLAIR | UnSup. Detection | 13.5 | 20.0 | 20.2 |
|  |  |  |  | 12.0 | 40.0 | 9.6 |
|  |  |  |  | 8.5 | 60.0 | 5.4 |
|  |  |  |  | 6.2 | 80.0 | 3.5 |

The performance summary of some of the existing works on 15 training images of MSSEG 2016 is presented in Table 4. The table includes both supervised^44,59–69^ and unsupervised^58,70,71^ methods. In the MSSEG 2016 challenge, the lesion detection capability was evaluated using various performance metrics, among which we have chosen to compare the Dice coefficient (DICE), TPR, and PPV^6^ in Table 4. From the table, we can see that different groups explored both multi-channel and single FLAIR MRI for processing. Beaumont et al.^58^ did not consider lesions appearing at the edges of the brain mask, not dominantly located in the WM, etc., for final refinement of lesion boundaries. In their following work, Beaumont et al.^70^ reported augmented accuracy after implementing post-processing technique. Knight and Khademi^71^ showed 3D Gaussian filtering with 0.5 SD to be efficient for noise removal in FLAIR. In this work, we also used the Gaussian filter, however with a higher SD (2.0) in computing local image statistics. They also used three false-positive reduction strategies based on lesion volume, distance from the edge of the brain, and distance from midline of the brain. In other works^59,60^, handcrafted features and post-processing were used for MS lesion segmentation. Most groups favored working with multi-contrast MRI^6^ for either designing machine learning algorithms or obtaining tissue prior, and followed with rule-based or morphological-operator-driven post-processing. Table 4 also includes the performances of the proposed and the baseline FFT detection approaches, similarly to MSLS 2008. The results on MSSEG 2016 show the same trend, i.e. the performance of the 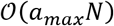 and FFT methods are very close, and the 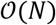 method achieves slightly better accuracy. In general, the sensitivity (TPR) of lesion detection seems moderate with a high false detection in both databases. The lower DICE in the three detection methods that we tested (compared to segmentation methods in the literature) is mainly attributed to the facts that: 1) we used a single image (as opposed to multiple images with different contrasts), 2) we performed no post-processing, 3) we employed no supervision, but most importantly 4) our goal was to detect (find the rough location of) as opposed to segment (delineate) the lesion. As such, the segmentation performance metrics used for the three detection methods should be compared among themselves, but not directly to those of the segmentation methods. The organizers of MSSEG 2016, Commowick et al.^6^, summarized that the deep-learning based approaches showed promise in narrowing the gap between manual and automatic segmentation of MS lesions. As our proposed method is a fully automatic unsupervised detection technique, the comparison with machine learning (deep neural networks) is not so informative due to the fundamental differences with supervised methods.

In summary, the proposed template matching algorithms do not require tissue prior or any lesion-specific information. They are fully automatic and unsupervised models that can detect lesions from the FLAIR sequence with a lower computational complexity than the conventional FFT-based template matching. We also did not apply any post-refinement approaches to remove the false positives; however, false lesions can be ruled out during the review by the clinician. In lesion detection, we give a higher priority to avoiding false negatives than false positives.

## Conclusions

We have proposed a new mathematical framework to compute a radius-optimized normalized cross-correlation coefficient (NCCC) similarity measure for 3D template matching, with application for automatic lesion detection. The key feature of our method is its low *linear-time* computational complexity, allowing for fast detection. Our alternative way to compute the NCCC exploits the approximation of Gaussian smoothing with multiple convolutions with a box kernel. We further proposed an analytical solution for the template radius that replaces the costly exhaustive search, thereby achieving the computational complexity of 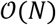, which is not possible with the FFT-based method. The proposed technique provides a new approach to implementing fast template matching for object detection from images. We tested our lesion-detection method on synthetic images as well as two real MRI databases of multiple sclerosis lesions, and compared its performance with existing state-of-the-art methods and the conventional FFT-based template matching algorithm. Future work includes extending our approach for anisotropic-voxel images, and also imaging modalities beyond MRI.

## Data Availability

The proposed methods were implemented on MATLAB platform. The codes can be made available upon request. The datasets used in this study for evaluating the proposed methods are available to public from http://www.ia.unc.edu/MSseg/index.html (MICCAI MSLS 2008) and https://portal.fli-iam.irisa.fr/msseg-challenge/data (MICCAI MSSEG 2016).

## Acknowledgements

Support for this research was provided in part by a Fulbright-Nehru Doctoral Research Fellowship (Award no: 2098/DR/2015-2016) and Research Associateship grant funded by Council of Scientific and Industrial Research, Govt. of India (File no. 09/81(1326)/18, EMR-1). IA was supported by the National Institutes of Health (NIH), specifically the National Institute of Diabetes and Digestive and Kidney Diseases (K01DK101631) and the National Institute on Aging (R56AG068261), and the BrightFocus Foundation (A2016172S). Computational resources were provided by the NIH Shared Instrumentation Grants (S10RR023401, S10RR019307, and S10RR023043).

## Author Contributions

S.K. developed the methods, performed the experiments, and wrote the manuscript. P.K.D. and I.A. supervised the work, reviewed the manuscript multiple times, and provided substantial feedback.

## References

1 Clinic, C. Brain Lesions, <https://my.clevelandclinic.org/health/diseases/17839-brain-lesions> (2019).

2 WebMD. Brain Lesions: Causes, Symptoms, Treatments, <https://www.webmd.com/brain/brain-lesions-causes-symptoms-treatments#1> (2019).

3 Egger, C. et al. MRI FLAIR lesion segmentation in multiple sclerosis: Does automated segmentation hold up with manual annotation? NeuroImage: Clinical 13, 264–270 (2017).

4 Schmidt, P. et al. An automated tool for detection of FLAIR-hyperintense white-matter lesions in multiple sclerosis. Neuroimage 59, 3774–3783 (2012).

5 Schmidt, P. Bayesian inference for structured additive regression models for large-scale problems with applications to medical imaging, PhD dissertation, IMU (2017).

6 Commowick, O. et al. Objective evaluation of multiple sclerosis lesion segmentation using a data management and processing infrastructure. Scientific reports 8, 1–17 (2018).

7 Lu, S., Lu, Z. & Zhang, Y.-D. Pathological brain detection based on AlexNet and transfer learning. Journal of computational science 30, 41–47 (2019).

8 Lu, S., Wang, S.-H. & Zhang, Y.-D. Detection of abnormal brain in MRI via improved AlexNet and ELM optimized by chaotic bat algorithm. Neural Computing and Applications, 1–13 (2020).

9 Cabezas, M. et al. Automatic multiple sclerosis lesion detection in brain MRI by FLAIR thresholding. Computer methods and programs in biomedicine 115, 147–161 (2014).

10 Ambrosini, R. D., Wang, P. & O’Dell, W. G. Computer-Aided Detection of Metastatic Brain Tumors Using Automated 3-D Template Matching. J Magn Reson Imaging 31, 85–93, doi:10.1002/jmri.22009 (2010).

11 Narasimha, R. et al. Evaluation of denoising algorithms for biological electron tomography. Journal of structural biology 164, 7–17 (2008).

12 Koley, S., Chakraborty, C., Mainero, C., Fischl, B. & Aganj, I. in International Workshop on Brainlesion: Glioma, Multiple Sclerosis, Stroke and Traumatic Brain Injuries. 52–61 (Springer).

13 Tourassi, G. D., Vargas‐Voracek, R., Catarious, D. M. & Floyd, C. E. Computer‐assisted detection of mammographic masses: A template matching scheme based on mutual information. Medical physics 30, 2123–2130 (2003).

14 Lochanambal, K., Karnan, M. & Sivakumar, R. in 2010 Second International Conference on Communication Software and Networks. 339–342 (IEEE).

15 Farag, A. A., El-Baz, A., Gimel’farb, G. & Falk, R. in Proceedings of the 17th International Conference on Pattern Recognition, 2004. ICPR 2004. 738–741 (IEEE).

16 Osman, O., Ozekes, S. & Ucan, O. N. Lung nodule diagnosis using 3D template matching. Computers in Biology and Medicine 37, 1167–1172 (2007).

17 Wang, P., DeNunzio, A., Okunieff, P. & O’Dell, W. G. Lung metastases detection in CT images using 3D template matching. Medical Physics 34, 915–922 (2007).

18 Moltz, J. H., Schwier, M. & Peitgen, H.-O. in 2009 IEEE International Symposium on Biomedical Imaging: From Nano to Macro. 843–846 (IEEE).

19 Warfield, S. K., Kaus, M., Jolesz, F. A. & Kikinis, R. Adaptive, template moderated, spatially varying statistical classification. Medical image analysis 4, 43–55 (2000).

20 Farjam, R., Parmar, H. A., Noll, D. C., Tsien, C. I. & Cao, Y. An approach for computer-aided detection of brain metastases in post-Gd T1-W MRI. Magnetic resonance imaging 30, 824–836 (2012).

21 Yang, S. et al. Computer-aided detection of metastatic brain tumors using magnetic resonance black-blood imaging. Investigative radiology 48, 113–119 (2013).

22 Wang, X.-F., Gong, J., Bu, R.-R. & Nie, S.-D. in Life System Modeling and Simulation. 50–61 (Springer, 2014).

23 Muñoz, A., Ertlé, R. & Unser, M. Continuous wavelet transform with arbitrary scales and O (N) complexity. Signal processing 82, 749–757 (2002).

24 Hou, H. & Andrews, H. Cubic splines for image interpolation and digital filtering. IEEE Transactions on acoustics, speech, and signal processing 26, 508–517 (1978).

25 http://mathworld.wolfram.com/SincFunction.html. Sinc Function. 2016.

26 Kogan, S. A note on definite integrals involving trigonometric functions. Available in http://www.mathsoft.com/asolve/constant/pi/sin/sin.html (1999).

27 Cooley, J. W. & Tukey, J. W. An Algorithm for the Machine Calculation of Complex Fourier Series. Mathematics of Computation 19, 297–301, doi:10.2307/2003354 (1965).

28 Johnston, B., Atkins, M. S., Mackiewich, B. & Anderson, M. Segmentation of multiple sclerosis lesions in intensity corrected multispectral MRI. IEEE transactions on medical imaging 15, 154–169 (1996).

29 Styner, M. et al. 3D segmentation in the clinic: A grand challenge II: MS lesion segmentation. Midas Journal 2008, 1–6 (2008).

30 Edlow, B. L. et al. 7 Tesla MRI of the ex vivo human brain at 100 micron resolution. BioRxiv, 649822 (2019).

31 Barkhof, F. & Scheltens, P. Imaging of white matter lesions. Cerebrovascular Diseases 13, 21–30 (2002).

32 Ruz, G. A., Estevez, P. A. & Perez, C. A. A neurofuzzy color image segmentation method for wood surface defect detection. Forest products journal 55, 52–58 (2005).

33 Fletcher, R. H., Fletcher, S. W. & Fletcher, G. S. Clinical epidemiology: the essentials. (Lippincott Williams & Wilkins, 2012).

34 Geremia, E. et al. in International Conference on Medical Image Computing and Computer-Assisted Intervention. 111–118 (Springer).

35 Brosch, T. et al. in International Conference on Medical Image Computing and Computer-Assisted Intervention. 3–11 (Springer).

36 Manjón, J. V. et al. MRI white matter lesion segmentation using an ensemble of neural networks and overcomplete patch-based voting. Computerized Medical Imaging and Graphics 69, 43–51 (2018).

37 Souplet, J.-C., Lebrun, C., Ayache, N. & Malandain, G. in MICCAI-Multiple Sclerosis Lesion Segmentation Challenge Workshop (New York, NY, USA, 2008).

38 Weiss, N., Rueckert, D. & Rao, A. in International Conference on Medical Image Computing and Computer-Assisted Intervention. 735–742 (Springer).

39 Wang, R. et al. Automatic segmentation and volumetric quantification of white matter hyperintensities on fluid-attenuated inversion recovery images using the extreme value distribution. Neuroradiology 57, 307–320 (2015).

40 Van Leemput, K., Maes, F., Vandermeulen, D., Colchester, A. & Suetens, P. Automated segmentation of multiple sclerosis lesions by model outlier detection. IEEE transactions on medical imaging 20, 677–688 (2001).

41 Prastawa, M. & Gerig, G. Automatic MS lesion segmentation by outlier detection and information theoretic region partitioning. Grand Challenge Work.: Mult. Scler. Lesion Segm. Challenge, 1–8 (2008).

42 Jerman, T., Galimzianova, A., Pernuš, F., Likar, B. & Špiclin, Ž. in BrainLes 2015. 45–56 (Springer).

43 Jesson, A. & Arbel, T. Hierarchical MRF and random forest segmentation of MS lesions and healthy tissues in brain MRI. Proceedings of the 2015 longitudinal multiple sclerosis lesion segmentation challenge, 1–2 (2015).

44 Valverde, S. et al. Improving automated multiple sclerosis lesion segmentation with a cascaded 3D convolutional neural network approach. NeuroImage 155, 159–168 (2017).

45 Ghribi, O. et al. An advanced MRI multi-modalities segmentation methodology dedicated to multiple sclerosis lesions exploration and differentiation. IEEE transactions on nanobioscience 16, 656–665 (2017).

46 Roura, E. et al. A toolbox for multiple sclerosis lesion segmentation. Neuroradiology 57, 1031–1043 (2015).

47 Geremia, E. et al. Spatial decision forests for MS lesion segmentation in multi-channel magnetic resonance images. NeuroImage 57, 378–390 (2011).

48 Guizard, N. et al. Rotation-invariant multi-contrast non-local means for MS lesion segmentation. NeuroImage: Clinical 8, 376–389 (2015).

49 Brosch, T. et al. Deep 3D convolutional encoder networks with shortcuts for multiscale feature integration applied to multiple sclerosis lesion segmentation. IEEE transactions on medical imaging 35, 1229–1239 (2016).

50 Strumia, M. et al. White matter MS-lesion segmentation using a geometric brain model. IEEE transactions on medical imaging 35, 1636–1646 (2016).

51 Zhan, T. et al. Multimodal spatial-based segmentation framework for white matter lesions in multi-sequence magnetic resonance images. Biomedical Signal Processing and Control 31, 52–62 (2017).

52 Anbeek, P., Vincken, K. L., Van Osch, M. J., Bisschops, R. H. & Van Der Grond, J. Probabilistic segmentation of white matter lesions in MR imaging. NeuroImage 21, 1037–1044 (2004).

53 Sudre, C. H. et al. Bayesian model selection for pathological neuroimaging data applied to white matter lesion segmentation. IEEE transactions on medical imaging 34, 2079–2102 (2015).

54 Tomas-Fernandez, X. & Warfield, S. K. A model of population and subject (MOPS) intensities with application to multiple sclerosis lesion segmentation. IEEE transactions on medical imaging 34, 1349–1361 (2015).

55 Bricq, S., Collet, C. & Armspach, J.-P. Unifying framework for multimodal brain MRI segmentation based on Hidden Markov Chains. Medical image analysis 12, 639–652 (2008).

56 Ong, K. H., Ramachandram, D., Mandava, R. & Shuaib, I. L. Automatic white matter lesion segmentation using an adaptive outlier detection method. Magnetic resonance imaging 30, 807–823 (2012).

57 Samaille, T., Colliot, O., Dormont, D. & Chupin, M. in 2011 IEEE international symposium on biomedical imaging: From nano to macro. 2014–2017 (IEEE).

58 Beaumont, J., Commowick, O. & Barillot, C. Multiple Sclerosis lesion segmentation using an automated multimodal Graph Cut. Proceedings of the 1st MICCAI Challenge on Multiple Sclerosis Lesions Segmentation Challenge Using a Data Management and Processing Infrastructure-MICCAI-MSSEG, 1–7 (2016)

59 Mahbod, A., Wang, C. & Smedby, O. Automatic multiple sclerosis lesion segmentation using hybrid artificial neural networks. Proceedings of the 1st MICCAI Challenge on Multiple Sclerosis Lesions Segmentation Challenge Using a Data Management and Processing Infrastructure-MICCAIMSSEG, 29–36 (2016).

60 Vera-Olmos, F., Melero, H. & Malpica, N. Random forest for multiple sclerosis lesion segmentation. Proceedings of the 1st MICCAI Challenge on Multiple Sclerosis Lesions Segmentation Challenge Using a Data Management and Processing Infrastructure-MICCAI-MSSEG, 81–86 (2016).

61 Salehi, S. S. M., Erdogmus, D. & Gholipour, A. in International workshop on machine learning in medical imaging. 379–387 (Springer).

62 Hashemi, S. R. et al. Asymmetric loss functions and deep densely-connected networks for highly-imbalanced medical image segmentation: Application to multiple sclerosis lesion detection. IEEE Access 7, 1721–1735 (2018).

63 Coupé, P., Tourdias, T., Linck, P., Romero, J. E. & Manjón, J. V. in International Workshop on Patch-based Techniques in Medical Imaging. 95–103 (Springer).

64 Chen, Z., Wang, X. & Zheng, J. in 2020 16th International Conference on Control, Automation, Robotics and Vision (ICARCV). 678–683 (IEEE).

65 Kamraoui, R. A. et al. Towards broader generalization of deep learning methods for multiple sclerosis lesion segmentation. arXiv preprint arXiv:2012.07950 (2020).

66 Valverde, S. et al. One-shot domain adaptation in multiple sclerosis lesion segmentation using convolutional neural networks. NeuroImage: Clinical 21, 101638 (2019).

67 Zhang, H. et al. in International Conference on Medical Image Computing and Computer-Assisted Intervention. 338–346 (Springer).

68 McKinley, R. et al. Simultaneous lesion and brain segmentation in multiple sclerosis using deep neural networks. Scientific reports 11, 1–11 (2021).

69 Isensee, F., Petersen, J., Kohl, S. A., Jäger, P. F. & Maier-Hein, K. H. nnu-net: Breaking the spell on successful medical image segmentation. arXiv preprint arXiv:1904.08128 1, 1–8 (2019).

70 Beaumont, J., Commowick, O. & Barillot, C. Automatic Muliple Sclerosis lesion segmentation from Intensity-Normalized multi-channel MRI. Proceedings of the 1st MICCAI Challenge on Multiple Sclerosis Lesions Segmentation Challenge Using a Data Management and Processing Infrastructure-MICCAI-MSSEG, 9–15 (2016)

71 Knight, J. & Khademi, A. MS Lesion Segmentation Using FLAIR MRI Only. Proceedings of the 1st MICCAI Challenge on Multiple Sclerosis Lesions Segmentation Challenge Using a Data Management and Processing Infrastructure-MICCAI-MSSEG, 21–28 (2016).

